# Resistance Exercise and Mechanical Overload Upregulate Vimentin for Skeletal Muscle Remodeling

**DOI:** 10.1101/2024.06.03.597241

**Authors:** Joshua S. Godwin, J. Max Michel, Cleiton A. Libardi, Andreas N. Kavazis, Christopher S. Fry, Andrew D. Frugé, Mariah McCashland, Ivan J. Vechetti, John J. McCarthy, C. Brooks Mobley, Michael D. Roberts

**Author notes:** Co-correspondence: Michael D. Roberts, PhD, Auburn University Alumni Professor, Nutrabolt Applied and Molecular Physiology Laboratory, School of Kinesiology, Auburn University, C. Brooks Mobley, PhD, Assistant Clinical Professor, Nutrabolt Applied and Molecular Physiology Laboratory, School of Kinesiology, Auburn University.

## Abstract

Our laboratory has performed various experiments examining the proteomic alterations that occur with mechanical overload (MOV)-induced skeletal muscle hypertrophy. In the current study we first sought to determine how 10 weeks of resistance training in 15 college-aged females affected protein concentrations in different tissue fractions. Training, which promoted significantly lower body muscle- and fiber-level hypertrophy, notably increased sarcolemmal/membrane protein content (+10.1%, p<0.05). Sarcolemmal/membrane protein isolates were queried using mass spectrometry-based proteomics, ∼10% (38/387) of proteins associated with the sarcolemma were up-regulated (>1.5-fold, p<0.05), and one of these targets (the intermediate filament vimentin; VIM) warranted further mechanistic investigation. VIM expression was first examined in the plantaris muscles of 4-month-old C57BL/6J mice following 10- and 20-days of MOV via synergist ablation. Relative to Sham (control) mice, VIM mRNA and protein content was significantly higher in MOV mice and immunohistochemistry indicated that VIM was predominantly present in the extracellular matrix (ECM). The 10- and 20-day MOV experiments were replicated in Pax7-DTA (tamoxifen-induced, satellite cell depleted) mice, which reduced the presence of VIM in the ECM. Finally, a third set of 10- and 20-day MOV experiments were performed in C57BL/6 mice intramuscularly injected with either AAV9-scrambled (control) or AAV9-VIM shRNA. While VIM shRNA mice presented with lower VIM in the ECM (∼50%), plantaris masses in response to MOV were similar between the injection groups. However, VIM shRNA mice presented with appreciably more MyHC_emb_-positive fibers with centrally located nuclei, indicating a regenerative phenotype. Using an integrative approach, we propose that skeletal muscle VIM is a mechanosensitive target predominantly localized to the ECM, and satellite cells are involved in its expression. Moreover, a disruption in VIM expression during MOV leads to dysfunctional skeletal muscle hypertrophy.

## INTRODUCTION

Skeletal muscle is a highly dynamic tissue that can undergo marked changes such as an increase in size (i.e., hypertrophy). The most potent stimulus for skeletal muscle hypertrophy is mechanical overload, either through progressive resistance training in humans or synergist ablation in rodents [1]. Mechano-sensitive protein signaling pathways are responsible for relaying the mechanical perturbations and tension placed on skeletal muscle into downstream signals for adaptations to occur [2–4]. A major pathway for mechanotransduction involves transmembrane proteins (integrins) and various accessory proteins that interact with the extracellular matrix (ECM), cytoskeletal network, and sarcolemma to propagate signals that enhance muscle protein synthesis [5–7].

Cytoskeletal and intermediate filaments not only provide structural support to the cell, but are also involved in processes like signal transmission, mechanotransduction, and gene regulation [8–10]. Intermediate filaments in muscle cells have also been linked to development [11, 12], regeneration [13, 14], and growth [15]. Desmin is a commonly studied muscle-specific gene that has been implicated in several processes that govern skeletal muscle homeostasis and is responsive to mechanical loading [16–18]. In contrast, the intermediate filament vimentin (VIM) is predominately expressed in immature developing myofibers and decreases in mature myofibers [19–21]. Interestingly, increased VIM expression is linked to active and proliferating satellite cells [19, 22] and appears to be critical for myofiber regeneration [23, 24]. VIM also plays a key role in several signaling pathways including mTORC1 [10]. Most notably, while not in skeletal muscle, a recent study reported that VIM-knockout cells had a reduction in mTORC1 signaling, blunted protein synthesis, and affected cell size [25]. However, the association of VIM with skeletal muscle hypertrophy has not been investigated, and its role in hypertrophy remains to be established.

Taken together, skeletal muscle hypertrophy in response to mechanical loading is dependent on several tightly regulated pathways and it appears that various ECM, cytoskeletal and intermediate filament-related proteins play important roles in this process. Furthermore, VIM has been shown to regulate cell size and protein synthesis response in other cell types. Herein, we performed shotgun proteomics on the sarcolemmal/ECM fraction of human skeletal muscle and show that: i) 10 weeks of resistance training increases VIM protein expression, and ii) a significant positive correlation exists between training-induced VIM protein levels and changes in mean muscle fiber cross-sectional area. Through a series of experiments in mice subjected to mechanical overload, we show that VIM expression in the ECM is upregulated during synergist ablation-induced skeletal muscle hypertrophy, and this response is dampened in mice lacking satellite cells. Finally, viral-mediated knockdown experiments in mice subjected to synergist ablation show that a blunting of VIM expression does not affect gross measures of hypertrophy in skeletal muscle but may delay/impair the regeneration and remodeling processes. Collectively, these data support that the upregulation of VIM occurs with mechanical overload-induced skeletal muscle hypertrophy, this process is partially dependent upon satellite cells, and a sufficient upregulation of VIM is needed to support proper skeletal muscle hypertrophy.

## METHODS

### Human Participants

Human muscle was from a study that was approved by the Auburn University Institutional Review Board (IRB) (Protocol # 19-249 MDR 1907), conformed to the standards set by the latest revisions of the Declaration of Helsinki, and was registered as a clinical trial (NCT04707963). The intent of the trial was to determine whether daily peanut protein supplementation (SUP), or no supplementation (control group [CON]), affected resistance training-related adaptations [26]. In the present study we utilized the remaining muscle samples, which included nine women from the SUP group and six women from the CON group. Our sub-sample showed no differences between SUP and CON mean fiber cross-sectional area (fCSA), *P* = 0.264), vastus lateralis cross-sectional area (by ultrasonography) (*P* = 0.421), thigh cross-sectional area (by peripheral computed tomography) (*P* = 0.855), as previously demonstrated [26]. Therefore, we combined data from the SUP and CON groups to perform the analyses presented herein. The samples were from 15 females (age: 21 ± 2 years, body mass index: 24.6 ± 3.7 kg/m^2^) who had not participated in resistance training programs, who were free of obvious cardiovascular or metabolic disease or other conditions contraindicating participation in exercise programs or donating muscle biopsies, and who were not pregnant or trying to become pregnant.

### Experimental Design for Human Training Study

Baseline assessments included muscle imaging assessments and vastus lateralis biopsies obtained after an overnight fast. Immediately afterwards, participants engaged in familiarization sessions with the RT protocol and test procedures. Approximately 3-5 days following familiarization sessions, the participants initiated an RT program with 2 sessions per week for 10 weeks. Seventy-two hours after the last RT session all assessments (including the procurement of a muscle biopsy) were repeated. For more details related to the study design and resistance training program, readers are referred to Sexton et al. [26].

### Vastus Lateralis Muscle Biopsies

Pre- and post-intervention vastus lateralis muscle biopsies were performed from the right leg. Initially the participants laid down on an athletic table where the upper thigh was shaved and cleaned with 70% isopropanol. A subcutaneous injection of 1% lidocaine (0.8 mL) was administered. After 5 minutes, the area was cleaned with chlorhexidine solution. Thereafter, a 7 mm-width pilot incision was made through the dermis with a sterile No. 11 surgical blade (AD Surgical; Sunnyvale, CA, USA). A sterile 5-gauge biopsy needle was then inserted into the pilot incision, through the muscle fascia, and ∼2 cm into the muscle (for a total depth of ∼4-7 cm) where a 40-80 mg sample was collected while applying suction. Following biopsies, tissue was rapidly teased of blood and connective tissue. A portion of the tissue (∼10–20mg) was preserved in optimal cutting temperature (OCT) media for histology (Tissue-Tek®, Sakura Finetek Inc.; Torrance, CA, USA), slowly frozen in liquid nitrogen-cooled isopentane, and subsequently stored at −80°C. Another portion of the tissue (∼30–50mg) was placed in pre-labeled foils, flash frozen in liquid nitrogen, and subsequently stored at −80°C for other molecular analyses described below. Tissue triage procedures were performed over a 3-minute period.

### Resistance Training Program

The resistance training program was described in detail previously [26]. Briefly, resistance training bouts consisted of participants performing a 45° leg press, leg extension, hex-bar deadlifts, flat barbell bench press, and wide-grip cable pulldown exercises. The same exercise order was followed for all training sessions, and two day per week training sessions consisted of performing 4 sets of 10 repetitions for each exercise one day per week, and 5 sets of 6 repetitions for each exercise one day per week. Two-minute rest intervals between sets and exercises were allotted throughout the training intervention. The training load was progressively increased on a session-by-session basis using the rating of perceived exertion (RPE) scale of 1-10 (“really easy” to “really hard”) so that each set was performed close to concentric muscle failure.

### Sarcoplasmic and Sarcolemmal Protein Isolations

Isolation of sarcoplasmic and sarcolemmal protein were initially isolated from muscle tissue using a high-fidelity membrane protein extraction kit according to the manufacturer’s recommendations (Mem-PRE Plus Membrane Protein Extraction Kit; Thermo Fisher Scientific; Waltham, MA, USA). This process yielded solubilized sarcoplasmic and sarcolemmal protein isolates.

The protein concentrations of both fractions were determined in duplicate on the same day as protein isolations to minimize freeze-thaw effects using a commercially available assay (bicinchoninic acid [BCA] Protein Assay Kit; Thermo Fisher Scientific). Thereafter, the fractions were stored at -80°C until subsequent analyses. The duplicate coefficient of variation (CV) values for the sarcoplasmic and sarcolemmal proteins readings were 2.3% and 2.4%, respectively.

### Proteomic of Sarcolemmal Protein Isolates

Sarcolemmal protein isolates were shipped to a commercial vendor for proteomics using a nanoLC-MS/MS platform (Creative Proteomics; Shirley, NY, USA). Briefly, the samples were treated with 50 mM ammonium bicarbonate and the resultant solutions were transferred to Microcon devices YM-10 (Millipore; Burlington, MA, USA). The samples were then centrifuged at 12,000 *g* at 4°C for 10 minutes. Protein concentrates were again treated with 50 mM ammonium bicarbonate and centrifuged again. Concentrates were then reduced using 10 mM DTT at 56°C for 1 hour and alkylated with 20 mM indole-3-acetic acid at room temperature in dark for 1 hour. Following reduction reactions, and one wash with 50 mM ammonium bicarbonate and centrifugation (12,000g at 4°C for 10 minutes), concentrates were incubated with 100 μL of 50 mM ammonium bicarbonate and free trypsin (ratio of 1:50) at 37°C overnight. The samples were then centrifuged at 12,000 *g* at 4°C for 10 minutes, and 100 μL of 50 mM ammonium bicarbonate was added to Microcon devices and centrifuged (for a total of two washes). Peptides were then isolated, lyophilized to near dryness, and resuspend in 20 μL of 0.1% formic acid for LC-MS/MS analysis.

Nanoflow ultra-high-performance liquid chromatography (UHPLC) was performed using an Ultimate 3000 nano UHPLC system (Thermo Fisher Scientific). Associated hardware included a trapping column (PepMap C18, 100Å, 100 μm×2 cm, 5 μm) and analytical column (PepMap C18, 100Å, 75 μm × 50 cm, 2 μm). Mobile phase A consisted of 0.1% formic acid in water, and mobile phase B consisted of 0.1% formic acid in 80% acetonitrile. The UHPLC runs consisted of 1 µL injections, a linear gradient from 2-8% buffer B for 3 minutes, from 8-20% buffer B for 56 minutes, 20-40% buffer B in 37 minutes, then from 4-90% buffer B for 4 minutes. Back-end mass spectrometry (MS) scans were performed between 300-1,650 m/z at a resolution of 60,000 at 200 m/z. The automatic gain control target for the full scan was set to 3e^6^. MS/MS scans were operated in Top20 mode using the following settings: resolution 15,000 at 200 m/z; automatic gain control target 1e^5^; maximum injection time 19 milliseconds; normalized collision energy at 28%; isolation window of 1.4 Th; charge sate exclusion: unassigned, 1, >6; dynamic exclusion 30 seconds.

MS files were analyzed against the HUMAN protein database using Maxquant (v1.6.1.14). Search parameters were set as follows: i) the protein modifications were carbamidomethylation (C) (fixed), oxidation (M) (variable), ii) the enzyme specificity was set to trypsin, iii) the maximum number of missed cleavages was set to 2, iv) the precursor ion mass tolerance was set to 10 ppm, and v) the MS/MS tolerance was 0.6 Da. A total of 1866 total peptides were identified, and these proteins were further queried using the PANTHER classification system (pantherdb.org) to delineate sarcolemmal membrane-associated proteins. These peptides were then subjected to statistical analyses where up-regulated and down-regulated proteins from pre-to-post training were considered to exceed a ±1.5-fold threshold (p<0.05) as recommended by Levin [27]. Notably, all proteomic data are presented as relative spectra values normalized to total spectra per sample.

### Mice Experiments

Mouse experiments not involving AAV9 injections (male and female wild-type C57BL/6 and male Pax7-DTA sham and overload mice) were conducted in accordance with the institutional guidelines for the care and use of laboratory animals as approved by the Animal Care and Use Committee of the University of Kentucky (protocol #: 2008-0291). All mice were housed in a climate-controlled room and maintained on a 14:10 hour light-dark cycle, food and water consumption were allowed *ad libitum*. For satellite cell depleted male mice the *Pax7^CreER/+^; Rosa26^DTA/+^* strain, called Pax7-DTA throughout, was generated as previously described [28]. Both sets of mice were subjected to dual-leg synergist ablation or sham surgeries (see below for description).

Mouse experiments involving AAV9 injections were approved by the Animal Care and Use Committee of the University of Nebraska (protocol #: 2091). Again, mice were housed in a climate-controlled room and maintained on a 14:10 hour light-dark cycle, food and water consumption were allowed *ad libitum*. For AAV9 experiments, male wildtype C57BL/6 mice were anesthetized under isoflurane, and plantaris muscles were injected with 30 uL (5x10^9^ particles) of either AAV9 VIM shRNA (catalog #: AA09-MSE096513-AVE001-A00) or AAV9 scrambled-shRNA (catalog #: SHD05); both of which were commercially developed (GeneCopoeia; Rockville, MD, USA) and contained a CMV promoter and GFP reporter. Following 30-days of viral incubation, mice were subjected to unilateral synergist ablation or sham surgeries.

### Conditional Ablation of Satellite Cells

Adult (≥4 months of age) Pax7-DTA mice were administered an intraperitoneal injection of either a vehicle (15% ethanol in sunflower seed oil) or tamoxifen (2 mg/day) for 5 consecutive days. Following a 2-week washout period, vehicle and tamoxifen-treated mice were divided into sham or synergist ablation (10-day or 20-day) groups.

### Synergist Ablation Surgeries in Mice

Surgical removal of synergist muscles (part of the gastrocnemius and the entire soleus) to mechanically overload the plantaris muscle was performed as previously described [28]. Briefly, mice were anesthetized with 3% isoflurane (with 1.5 L of O_2_ per minute) and placed in sternal recumbence where a longitudinal incision was made on the dorsal aspect of the lower hindlimb, and the tendon of the gastrocnemius muscle was isolated and used as a guide to excise the soleus and part of the gastrocnemius. The incision was then sutured, and the animals were allowed to recover in their home cages. Sham surgeries involved similar incision and suture procedures without the excision of muscles as described above. Ten- or 20-days following surgeries, mice were anesthetized, euthanized via cervical dislocation, and the plantaris muscle was excised. Immediately following removal, the plantaris muscles were weighed and either flash-frozen in liquid nitrogen or prepared for immunohistochemical analysis via preservation in OCT media.

### Human Muscle Immunohistochemistry

*Human fCSA Determination.* Human skeletal muscle samples were examined for fCSA using immunohistochemistry (IHC) as previously described by our laboratory [26]. Briefly, slides with sections were dried for 10 minutes at room temperature. Triton-X (0.5%) in phosphate buffer solution (PBS) was used to permeabilize the sections for 5 minutes. Thereafter, slides were washed in PBS for 5 minutes and subsequently incubated in blocking solution for 15 minutes (Pierce Super Blocker; Thermo Fisher Scientific). The slides were incubated in primary antibody solution (1x PBS, 5% of Pierce Super Blocker Solution, and 2% [or a 1:50 dilution] of rabbit anti-dystrophin IgG1 [catalog #: GTX15277; Genetex Inc.; Irvine, CA, USA] and mouse anti-myosin I IgG1 (catalog #: A4.951 Developmental Studies Hybridoma Bank; Iowa City, IA, USA) for 1 hour. Slides were then washed for 5 minutes in PBS and incubated in a secondary antibody solution (1x PBS and 1% [or 1:100 dilution] Texas Red-conjugated anti-rabbit IgG [catalog #: TI-1000; Vector Laboratories, Burlingame, CA, USA] and Alexa Fluor 488-conjugated anti-mouse IgG1 (catalog #: A-11001; Thermo Fisher Scientific) for 1 hour in the dark at room temperature. The slides were then mounted in fluorescent media and imaged using a fluorescent microscope (Nikon Instruments, Melville, NY, USA) with a 10x objective lens. Open-sourced software (MyoVision) was used to analyze all images for mean fCSA [29].

*Phalloidin staining.* To determine myofibril area per fiber, F-actin labelling using phalloidin-conjugated to Alexa Fluor 594 (AF594) was performed. Briefly, serial sections were allowed to air dry for ∼1.5–2 hours at room temperature followed by fixation in chilled acetone for 5 minutes. Sections were then incubated with 3% H_2_O_2_ for 15 minutes and true black (catalog #: 23007 Biotium; Fremont, CA, USA) for 1 minute to reduce auto-fluorescence. Sections were then blocked with 2.5% BSA/5% goat serum for 1 hour. The sections were incubated overnight at 4°C with mouse anti-dystrophin MANDY S8 (1:20) (catalog #: 8H11; Developmental Studies Hybridoma Bank). The following morning the sections were incubated for 1 hour with phalloidin conjugated to AF594 (1:100) (catalog #: A12381; Thermo Fisher Scientific) and Alex Fluor 488-conjugated anti-mouse IgG1 (1:250) (catalog #: A11001; Thermo Fisher Scientific). Sections were incubated with DAPI (1:10,000) (catalog #: D3571; Thermo Fisher Scientific) for 15 minutes. Slides were then mounted with glass cover slips using 50/50 PBS+glyercol. Digital images were captured using a fluorescent microscope (Nikon Instruments) using a 20x objective. Quantification of myofibril area per fiber was performed using ImageJ software (National Institutes of Health; Bethesda, MD, USA) according to previous reports [30, 31]. Briefly, images were split into RGB channels, and the red channel image was converted to grayscale. The threshold function was then used to generate a binary black/white image of stained versus portions of fibers. The myofibers were then traced, and myofibril areas were calculated as a percentage per myofiber area. The coefficient of variation (CV) between two measurements performed 72 hours apart was 0.26% [31].

### Mouse Plantaris Immunohistochemistry

*VIM area and fCSA determinations.* Mouse plantaris muscles preserved for IHC were sectioned at a thickness of 7 μm using a cryotome (Leica Biosystems; Buffalo Grove, IL, USA) and adhered to multiple positively charged slides for the interrogation of various targets as explained in this and subsequent paragraphs. Following section adhesion to slides, slides were allowed to air dry prior to being stored at -80°C until further analysis.

Sections from sham mice and mice subjected to mechanical overload were first stained for VIM, dystrophin, and DAPI. Briefly, sections were fixed in ice cold acetone for 5 minutes followed by 3% hydrogen peroxide (H_2_O_2_) for 10 minutes and true black incubation for 1 minute to reduce autofluorescence. Sections were then blocked with 5% goat serum/2.5% BSA for at least 1 hour before overnight incubation at 4°C with an antibody cocktail consisting of rabbit anti-vimentin (1:100) (catalog #: 5741; Cell Signaling Technologies; Danvers, MA, USA), mouse anti-dystrophin IgG2b (1:50) (catalog ID: Mandy S8; Developmental Studies Hybridoma Bank), and 2.5% BSA. After primary antibody incubations, sections were incubated for 1 hour with a secondary antibody cocktail consisting of anti-rabbit AF488 (Vector Laboratories), anti-mouse IgG2b AF594 (catalog #: A-21145; Thermo Fisher Scientific), and PBS. Following secondary antibody incubations, sections were incubated with DAPI (1:10,000) (catalog #: D3571; Thermo Fisher Scientific) for 10 minutes. Coverslips were then added using PBS + glycerol as mounting media. Digital images of entire sections were captured using a fluorescent microscope (20x objective; Zeiss Axio imager.M2) and motorized scanning stage.

*Satellite Cell and VIM Co-expression.* Sections stained for satellite cells and VIM to determine co-expression were first fixed in ice cold acetone for 5 minutes followed by incubation with 3% H_2_O_2_ for 10 minutes. Sections were then subjected to an epitope retrieval protocol. This protocol first involved incubating sections with sodium citrate for 2 minutes at room temperature followed by incubation in pre-warmed (65°C) sodium citrate. Sections were then placed in a water bath incubator until they reached 92°C. Sections were then incubated for 11 minutes at 92°C, allowed to cool in water bath set to 50°C, and then cooled at room temperature. Following epitope retrieval, sections were incubated with true black for 1 minute to reduce autofluorescence. Sections were then blocked using 5% goat serum/2.5% BSA for at least 1 hour followed by the addition of biotin and streptavidin blocking solutions for 15 minutes each. Sections were incubated with an antibody cocktail containing: i) mouse anti-Pax7 IgG1 (1:100), ii) mouse anti-dystrophin IgG2b (1:50), rabbit anti-vimentin (1:100), and 2.5% BSA overnight at 4°C. Following primary antibody incubations, sections were incubated with biotin-SP-conjugated goat anti-mouse IgG1 (1:1,000) (catalog #: 111-065-003; Jackson ImmunoResearch; West Grove, PA, USA) in 2.5% BSA for 90 minutes. Sections were then incubated with a secondary antibody cocktail consisting of SA-HRP (1:500) (catalog #: S-911; Thermo Fisher Scientific), goat anti-rabbit AF488 (1:250) (Vector Laboratories), and goat anti-mouse IgG2b AF647 (1:250) (catalog #: A21242; Thermo Fisher Scientific) for one hour. TSA AF555 (1:200) (catalog #: B-40955; Thermo Fisher Scientific) was added for 20 minutes, followed by DAPI for 10 minutes. Coverslips were added using PBS + glycerol as mounting medium. Digital images of entire sections were captured using a fluorescent microscope (20x objective; Zeiss Axio imager.M2) and motorized scanning stage.

*Fibroblast and VIM Co-expression.* Sections stained for fibroblasts and vimentin to determine co-expression were first fixed with ice cold acetone for 5 minutes, followed by incubation with 3% H_2_O_2_ for 10 minutes. Sections then underwent epitope retrieval as described above, followed by incubation with true black for 1 minute to reduce autofluorescence. Sections were then blocked with 5% goat serum/2.5% BSA for at least one hour followed by blocking with biotin and streptavidin solution for 15 minutes each. Following the blocking step, sections were incubated overnight at 4°C with rabbit anti-TCF4 (1:100) (catalog #: 2569; Cell Signaling Technologies) in 2.5% BSA. Sections were then incubated with goat biotin-conjugated anti-rabbit IgG (1:1000) (catalog #: 111-065-046; Jackson ImmunoResearch) in 2.5% BSA for 90 minutes, followed by incubation with SA-HRP (1:500) for 1 hour. Sections were then incubated with TSA-488 (1:400) (catalog #: B-40953; Thermo Fisher Scientific) for 20 minutes followed by incubation with DAPI for 10 minutes. Sections were briefly checked under microscope to ensure proper staining for TCF4 before being re-fixed in 4% PFA for 10 minutes followed by blocking with 5% goat serum/2.5% BSA and biotin/streptavidin solutions for at least 1 hour. Sections were then incubated overnight at 4°C with antibody cocktail consisting of mouse anti-dystrophin IgG2b (1:50), rabbit anti-vimentin (1:100), and 2.5% BSA. Sections were then incubated with a secondary antibody cocktail consisting of goat anti-rabbit AF555 (1:250), and goat anti-mouse IgG2b AF647 (1:250) for 1 hour, followed by incubation with DAPI for 10 minutes. Coverslips were added using PBS + glycerol as mounting medium. Digital images of entire sections were captured using a fluorescent microscope (20x objective; Zeiss Axio imager.M2) and motorized scanning stage.

*Fibro-adipogenic Progenitor and VIM Co-expression*. Only wildtype mouse plantaris sections were stained for fibro-adipogenic progenitors (FAPs) and vimentin to determine the co-expression of the two proteins using IHC. Briefly, the sections were fixed with 4% PFA for 10 minutes, incubated with 3% H_2_O_2_ for 15 minutes, and incubated with true black solution for 1 minute to reduce autofluorescence. Sections were then blocked with 2.5% BSA/0.1% Triton-X for 1 hour and subsequently incubated with an antibody cocktail consisting of goat anti-mouse PDGFRα (1:100) (catalog #: AF1062; R&D Systems; Minneapolis, MN, USA), rabbit anti-vimentin (1:100), and 2.5% BSA overnight at 4°C. Sections were then incubated for two hours with a secondary antibody cocktail consisting of donkey anti-goat AF555 (1:250) (catalog #: A-21432; Thermo Fisher Scientific), donkey anti-rabbit AF488 (1:250) (catalog #: A-21206; Thermo Fisher Scientific), and Wheat germ agglutinin (WGA) (1:50) (catalog #: W-32466; Thermo Fisher Scientific). The sections were then incubated with DAPI for 10 minutes and coverslips were added using PBS + glycerol as mounting medium. Digital images of entire sections were captured using a fluorescent microscope (20x objective; Zeiss Axio imager.M2) and motorized scanning stage.

*Macrophage and VIM Co-expression.* Again, only wild type mouse muscle was stained for macrophages and VIM to determine the co-expression of the two using IHC. Briefly, sections were fixed with ice cold acetone for 10 minutes followed by incubated with 3% H_2_O_2_ for 10 minutes. Sections were then blocked with 5% goat serum/2.5% BSA for at least one hour. Sections were then incubated overnight at 4°C with antibody cocktail consisting of rat anti-F4/80 (1:100) (catalog #: MCA497; Bio-Rad Laboratories; Hercules, CA, USA), rabbit anti-vimentin (1:100), and mouse anti-dystrophin IgG2b (1:50). Following overnight incubation with primary antibodies, sections were incubated with goat anti-rat biotin (1:500) (catalog #: 31830; Thermo Fisher Scientific) in 2.5% BSA for 90 minutes. The sections were then incubated for one hour with a secondary antibody cocktail consisting of SA-HRP (1:500), goat anti-rabbit AF488 (1:250), and goat anti-mouse IgG2b AF647 (1:250). TSA AF555 (1:200) was added for 20 minutes, followed by DAPI for 10 minutes. Coverslips were added using PBS + glycerol as mounting medium. Digital images of entire sections were captured using a fluorescent microscope (20x objective; Zeiss Axio imager.M2) and motorized scanning stage.

*Embryonic Myosin Heavy Chain.* AAV9-shRNA mouse plantaris muscles were stained for the presence of embryonic myosin heavy chain (MyHC_emb_). Briefly, the sections were fixed with ice cold acetone for 5 minutes followed by an incubation with 3% H_2_O_2_ for 10 minutes. Sections were then blocked with 5% goat serum/2.5% BSA for at least one hour. Sections were then incubated overnight at 4°C with an antibody cocktail consisting of mouse anti-dystrophin IgG2b (1:50) and mouse anti-eMHC IgG1 (1:50) (catalog ID: F1.652; Developmental Studies Hybridoma Bank). The following day, sections were incubated with a secondary antibody cocktail for one hour consisting of goat anti-mouse IgG2b AF647 (1:250) and goat anti-mouse IgG1 AF488 (1:250). Sections were then incubated with DAPI for 10 minutes and coverslips were added using PBS + glycerol as mounting medium. Digital images of entire sections were captured using a fluorescent microscope (20x objective; Zeiss Axio imager.M2) and motorized scanning stage.

### Cell Culture Experiments

Low passage (passage 3-5) immortalized C2C12 myoblasts (ATCC; Manassas, VA, USA) were used to determine vimentin protein expression throughout various stages of proliferation and differentiation as well as in response to an anabolic stimulus (insulin-like growth factor, IGF-1), and experiments were performed in a humidified CO_2_ incubator at 37°C using 5% CO_2_-95% room air. Myoblasts (100,000 cells per mL) were seeded onto six-well plates in growth medium (GM) containing Dulbecco’s modified Eagle’s Medium (DMEM; Corning, Corning, NY) supplemented with 10% fetal bovine serum and 1% penicillin/streptomycin. Once cells reached confluency (∼85-90%), differentiation was induced by switching to differentiation medium (DM), which consisted of DMEM supplemented with 2% horse serum and 1% penicillin/streptomycin. Once complete differentiation into myotubes was achieved, cells in one experiment were treated with 10 μM arabinosylcytosine (AraC; Sigma, St. Louis, MO, USA) for 24 hours to reduce the number of unfused myoblasts [32]. The myotubes were then treated with either PBS (CTL) or various doses (100 ng/mL, 200 ng/mL, 400 ng/mL) of recombinant mouse IGF-1 (catalog #: 791-MG-050; R&D Systems) for 24 hours. A separate set of plates that did not receive AraC treatments was used to determine how the enrichment of unfused myoblasts affected VIM expression. Finally, a third set of plates was examined at 0 days, 1 day, 3 days, 5 days, and 7 days after the initiation of differentiation to determine how differentiation affects VIM expression. At the end of experimental time lines, cells were washed with PBS, 500 μL of 1× cell lysis buffer (catalog #: 9803S; Cell Signaling Technologies) was added to wells, and wells were scraped using rubber policemen to collect lysates. The lysates were then centrifuged at 500 *g* for 5 minutes (4°C), supernatants were transferred to 1.7 mL microtubes, and lysates were stored at -80°C until western blotting.

### Western Blotting and qPCR

Western blotting as performed as previously described by our laboratory [33] using the rabbit anti-vimentin antibody described above (1:1,000). For myotube lysate western blotting, protein concentrations were determined using a commercially available BCA kit (Thermo Fisher Scientific). Cell lysates were prepared at a concentration of 1 μg/μL and 15 μL of sample was loaded onto 4-15% gradient SDS-polyacrylamide gels (Bio-Rad Laboratories). Following the electrotransfer of proteins onto PVDF membranes, blocking, and antibody incubations, membranes were developed using a chemiluminescent substrate (EMD Millipore) and imaged using a gel documentation system (ChemiDoc Touch; Bio-Rad Laboratories). Vimentin protein band densities were obtained using associated software (Image Lab v6.0.1; Bio-Rad Laboratories) and normalized to Ponceau lane densities. Target/Ponceau density ratios were normalized to either vehicle-control treatments (IGF-1 experiments) or day 0 differentiation (differentiation phase experiments).

Plantaris muscle RNA and protein from one mouse experiment (Figure 2 in Results) were isolated using the modified Trizol-bromochloropropane protocol described by Wen et al. [34]. Following RNA isolation, the RNA pellet was resuspended in 30 μL of RNase-free water and RNA concentrations were determined in duplicate at an absorbance of 260 nm by using a NanoDrop Lite (Thermo Scientific). Thereafter, cDNA (1 μg) was synthesized using a commercial qScript cDNA SuperMix (Quanta Biosciences, Gaithersburg, MD, USA) per the manufacturer’s recommendations. qPCR was performed with gene-specific primers and SYBR-green-based methods (Quanta Biosciences) with gene-specific primers designed with primer design software (Primer3Plus, Cambridge, MA, USA) using a real-time PCR thermal cycler (Bio-Rad Laboratories). Protein was isolated from the organic phase as described by Wen et al. [34], and VIM was interrogated using the rabbit anti-vimentin antibody described above (1:1,000). VIM protein band densities were obtained as described in the previous paragraph and normalized to Ponceau lane densities. Target/Ponceau density ratios were normalized to the Sham group.

**Figure 1.**
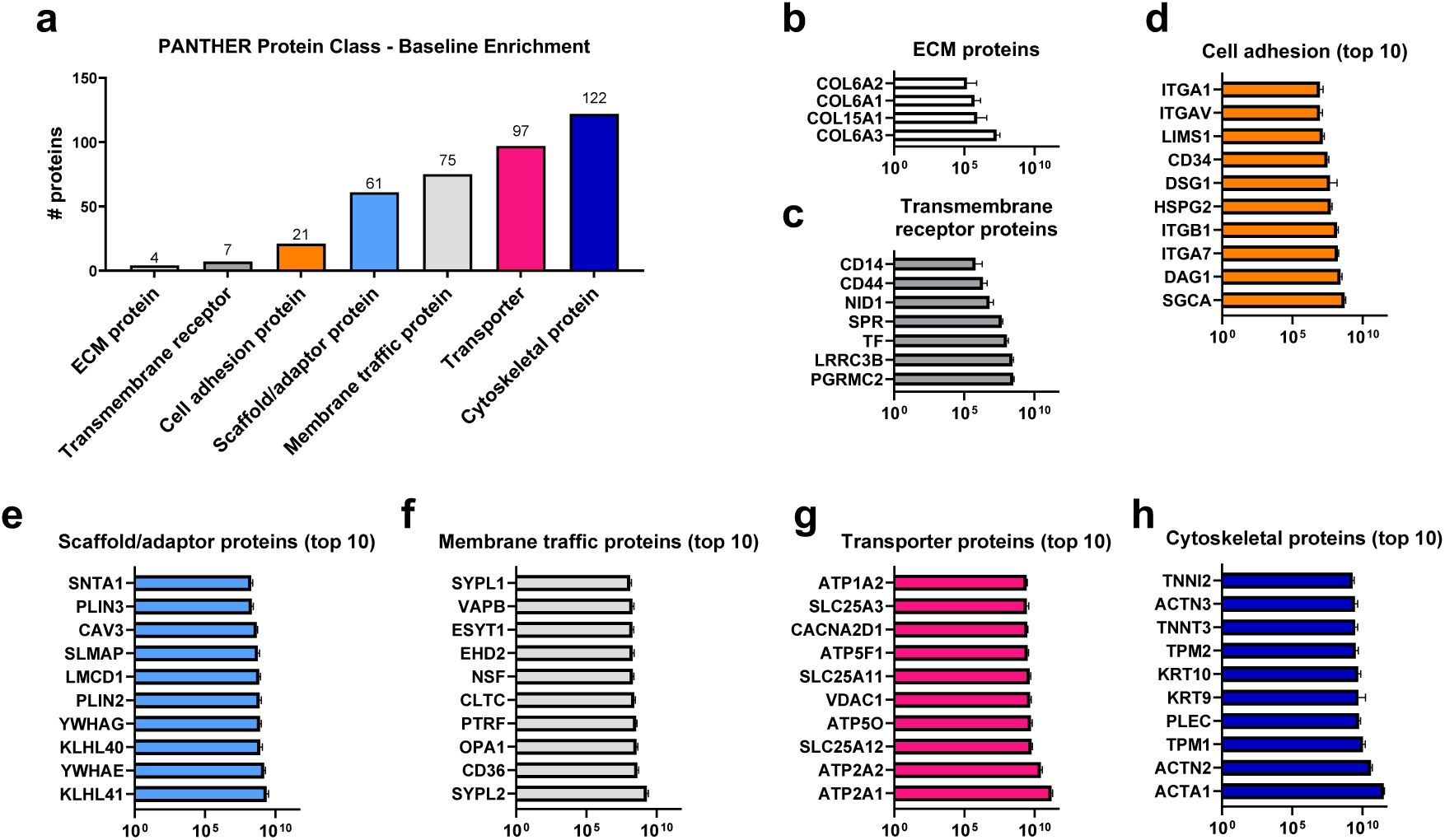
Basal state (pre-intervention) human sarcolemmal proteome data. a) represents the number of targets per PANTHER protein class identified in the sarcolemmal protein isolates via shotgun proteomics from pre-training human skeletal muscle biopsies (grand sum = 387 total targets identified). b-h) are the top enriched proteins (listed by gene name) for each protein class; note, d-h are limited to the top 10 targets due to space limitations. Additional note: bar graph data for b-h) are presented as mean and standard deviation values for all 15 participants. Raw data can be found in Supplemental File 2.

**Figure 2.**
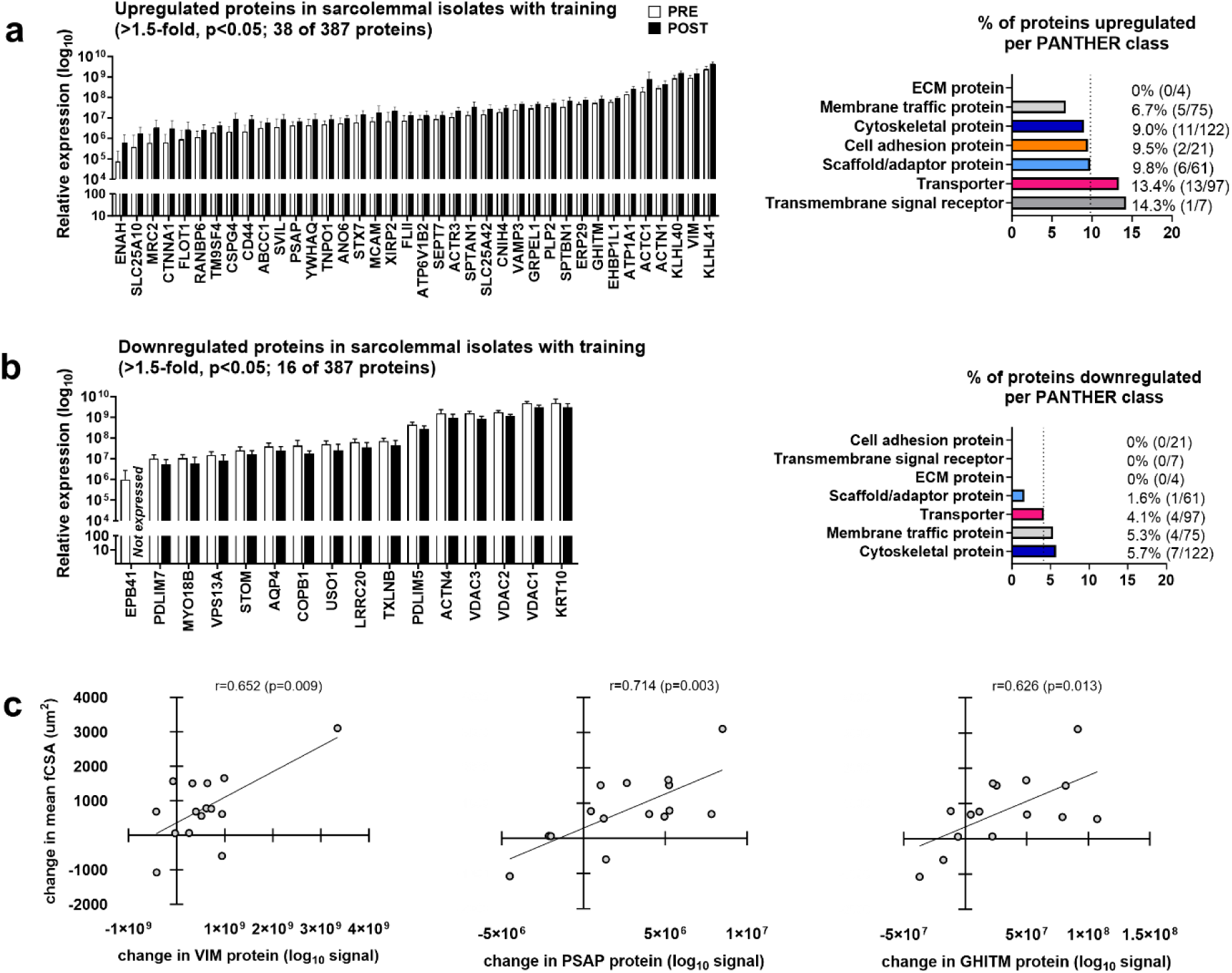
Human sarcolemmal proteome response to resistance training. a) upregulated proteins identified in the sarcolemmal fraction following training including percentage of proteins up-regulated upregulated in each PANTHER protein class. b) downregulated proteins identified in the sarcolemmal fraction following training including percentage of proteins down-regulated upregulated in each PANTHER protein class. c) significant correlations between pre-to-post changes in sarcolemmal proteins that were significantly up-regulated following resistance training and mean fiber cross-sectional area. Additional note: bar graph data for a/b) are presented as mean and standard deviation values for all 15 participants.

### Statistical Analyses

Statistical analyses were performed using GraphPad Prism (Version 10.2; San Diego, CA, USA). Human dependent variables (including proteomic data) were analyzed using dependent samples t-tests. All mouse experiments variables were analyzed using one-way ANOVAs and Tukey’s post-hoc tests were applied when significant model effects were observed. Pax7-DTA and AAV9 mouse experiment variables were analyzed using two-way ANOVAs with Tukey’s post-hoc tests being performed when significant main effects or interactions were observed. Select associations between dependent variables were also analyzed using Pearson correlations. All data are presented as mean ± standard deviation (SD) values, and a two-tailed statistical significance of p<0.05 was used throughout.

## RESULTS

### Pre-intervention Sarcolemmal Proteome Characteristics and Adaptions following 10 Weeks of Resistance Training

Given that the sarcolemmal proteome of human skeletal muscle has not been investigated to our knowledge, Figure 1 provides highlights of these data from the pre-intervention biopsies of the 15 females. Notably, 387 proteins were detected, and PANTHER protein classification analysis revealed the enrichment of 4 ECM proteins, 7 transmembrane signal receptor proteins, 21 cell adhesion proteins, 61 scaffold/adaptor proteins, 75 membrane traffic proteins, 97 transporter proteins, and 122 cytoskeletal proteins (Figure 1a). The top enriched proteins from each PANTHER class are also presented in Figure 1b-h.

Resistance training significantly increased whole-body lean mass (2.1%), mean (type I & II) fiber cross-sectional area (17.8%), and deadlift strength (32%) (Supplemental Figure 1). Sarcolemmal protein concentrations also significantly increased with 10 weeks of training (10.1%; Supplemental Figure 1). Resistance training upregulated 9.6% (38/387) of the detected proteins (>1.5-fold increase, p<0.05; Figure 2a), and PANTHER protein classes shown to be upregulated above this 9.6% threshold included scaffold/adaptor proteins (9.8%, 6/61), transporter proteins (13.4%, 13/97), and transmembrane signal receptor proteins (14.3%, 1/7). Training downregulated 4.1% (16/387) of the detected proteins (>1.5-fold decrease, p<0.05; Figure 2b), and PANTHER protein classes shown to be downregulated at or above this 4.1% threshold included transporter proteins (4.1%, 4/97), membrane traffic proteins (5.3%, 4/75), and cytoskeletal proteins (5.7%, 7/122; Figure 2b).

The pre-to-post intervention change scores of 3 enriched proteins upregulated with training were significantly correlated to changes in mean fiber cross sectional area (fCSA) including the cytoskeletal protein vimentin (VIM; r = 0.652, p = 0.009), the transporter protein prosaposin (PSAP; r = 0.714, p = 0.003), and transporter protein growth hormone-inducible transmembrane protein (GHITM; r = 0.626 p = 0.013) (Figure 2c). We decided to further pursue the role and function of VIM in the context of skeletal muscle hypertrophy for various reasons. First, there is compelling literature on VIM being linked to anabolic processes [25]. Second, the relative abundance of VIM was appreciably higher compared to the other two targets (see Figure 2a). Finally, we recently performed an independent eight-week unilateral leg resistance training study in middle-aged males and females [35]. Pre- and post-intervention muscle biopsies were analyzed using immunohistochemistry and deep proteomics. As with the current data, the training-induced change in VIM abundance was significantly correlated to changes in mean fiber cross sectional area (fCSA) (r = 0.514, p = 0.029, n=18 participants).

### Vimentin Expression Following 10- and 20-days Synergist Ablation in Mice

To further investigate the potential mechanistic role of VIM in muscle hypertrophy, we used mechanical overload (MOV) induced by synergist ablation in mice. Following 10- and 20-days of mechanical overload (MOV) via synergist ablation, VIM expression was measured at the mRNA level via qPCR, at the protein level via western blotting, and visually using immunohistochemistry. *Vim* mRNA was significantly upregulated with mechanical overload (ANOVA p < 0.001), and post hoc testing revealed that the 10-day MOV values were greater than Sham and 20-day MOV mice (p < 0.001, and p = 0.007, respectively), whereas values in 20-day MOV mice were not statistically different from Sham mice (p = 0.181; Figure 3a). Protein expression of VIM was also significantly upregulated with MOV (p < 0.001). Like mRNA expression patterns, VIM protein was upregulated in 10-day MOV compared Sham and 20-day MOV mice (p < 0.001 for both comparisons) and levels were not statistically different between Sham and 20-day MOV mice (p = 0.059; Figure 3b). Immunohistochemistry indicated that VIM was exclusively localized to the ECM (Figure 3f), and the percent area occupied by VIM in 20x objective fields of view was significantly higher in MOV versus Sham mice (ANOVA p < 0.001, 10-day versus Sham p < 0.001, 20-day versus Sham p = 0.006; Figure 3c). When Figure 3c data were normalized to fiber number, VIM content was still shown to be significantly higher in MOV versus Sham mice (ANOVA p < 0.001, 10-day versus Sham p < 0.001, 20-day versus Sham p = 0.006; Figure 3d). Kruskal-Wallis tests with Bonferroni correction indicate no difference in fCSA distribution between Sham and 10-day (p = 1.000), however, 20-day fCSA distribution was higher than Sham and 10-day (p < 0.001 for both; Figure 3e). Other muscle characteristics in response to 10-20 days of MOV are presented in Supplemental Figure 2.

**Figure 3.**
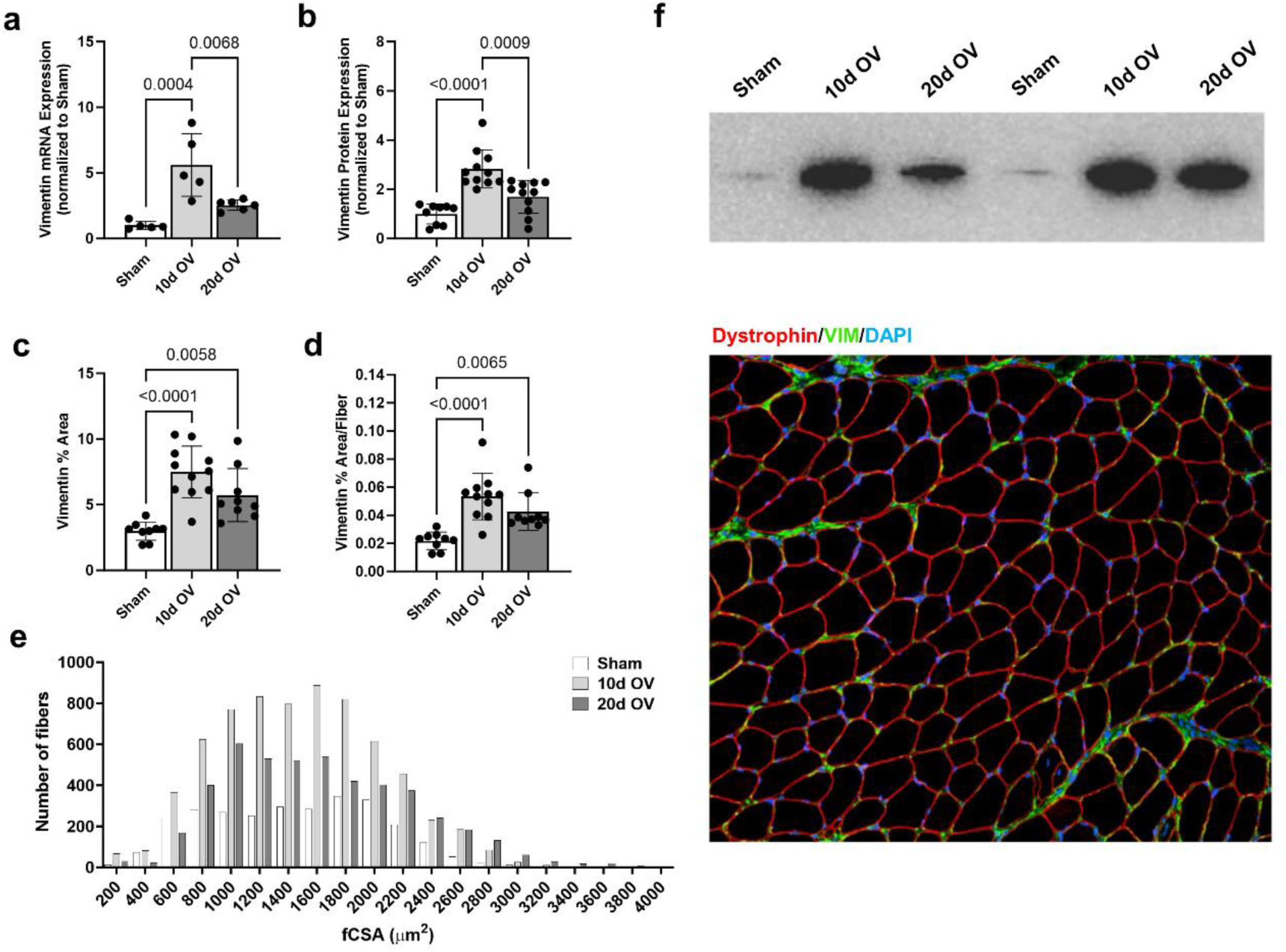
Plantaris muscle vimentin expression response to 10- and 20-days of mechanical overload via synergist ablation. All bar graph data (aside from histogram data) are presented as mean and standard deviation values with qPCR data containing 4-6 mice per condition and other data containing 9-11 mice per condition. a) mRNA expression of VIM via qPCR following 10- and 20-days MOV. b) Protein expression of VIM via western blotting following 10- and 20-days of MOV. c) Percent area of plantaris cross-section occupied by VIM following 10- and 20-days of MOV (determined via IHC). d) Percent area of plantaris cross-section occupied by VIM (normalized to fiber number) following 10- and 20-days of MOV. e) Fiber size distribution following 10- and 20-day MOV. f) Representative western blot and IHC images. Additional representative IHC images can be found in Supplemental Figure 6.

### Vimentin Co-expression with Resident Skeletal Muscle Stromal Cells

Given visual confirmation that VIM is localized to the ECM, we decided to measure the co-expression of this target with different stromal cells residing in the ECM during 10- and 20-day of MOV. We first queried the Tabula Muris single cell sequencing database [36] (method: FACS; tissue: limb muscle), and results indicated highest enrichments in macrophages and satellite cells. We then decided to investigate the co-expression of VIM with satellite cells, fibroblasts, fibro-adipocyte progenitor cells (FAPs), and macrophages in Sham as well as 10-day and 20-day MOV mice using immunohistochemistry. Satellite cells (Pax7+/DAPI+ cells per fiber) were significantly higher in MOV versus Sham mice (ANOVA p < 0.001, 10-day versus Sham p < 0.001, 20-day versus Sham p = 0.002; Figure 4a). VIM area (normalized to satellite cell number) was also significantly higher in MOV versus Sham mice (ANOVA p = 0.001, 10-day versus Sham p = 0.003, 20-day versus Sham p = 0.003; Figure 4a). Although fibroblasts (TCF4+/DAPI+ cells per fiber) were significantly higher in MOV versus Sham mice (ANOVA p < 0.001, 10-day versus Sham p < 0.001, 20-day versus Sham p = 0.003), VIM area (normalized to fibroblast number) was not affected with MOV (ANOVA p = 0.581; Figure 4b). While FAP counts (PDGFRα+/DAPI+ cells per fiber) were not affected with MOV (p = 0.280), VIM area (normalized to FAP number) was significantly higher in MOV versus Sham mice (ANOVA p < 0.001, 10-day versus Sham p < 0.001, 20-day versus Sham p < 0.001; Figure 4c). Macrophages (F4/80+/DAPI+ cells per fiber) were significantly higher in 10-day MOV versus Sham mice (ANOVA p < 0.001, 10-day versus Sham p < 0.001). However, values in 20-day versus 10-day MOV mice were lower (p = 0.016), and 20-day MOV versus Sham mice values were similar (p = 0.366; Figure 4d). VIM area (normalized to macrophage number) was significantly higher in MOV versus Sham mice (ANOVA p < 0.001, 10-day versus Sham p < 0.001, 20-day versus Sham p < 0.001; Figure 4d), albeit this marker was lower in 20-day versus 10-day mice (p = 0.024). Figure 4e-h shows representative images from these experiments. In sum, satellite cells were unique to other measured cell types given that satellite cell counts (and increased VIM protein per satellite cell) increased with 10- and 20-days of MOV.

**Figure 4.**
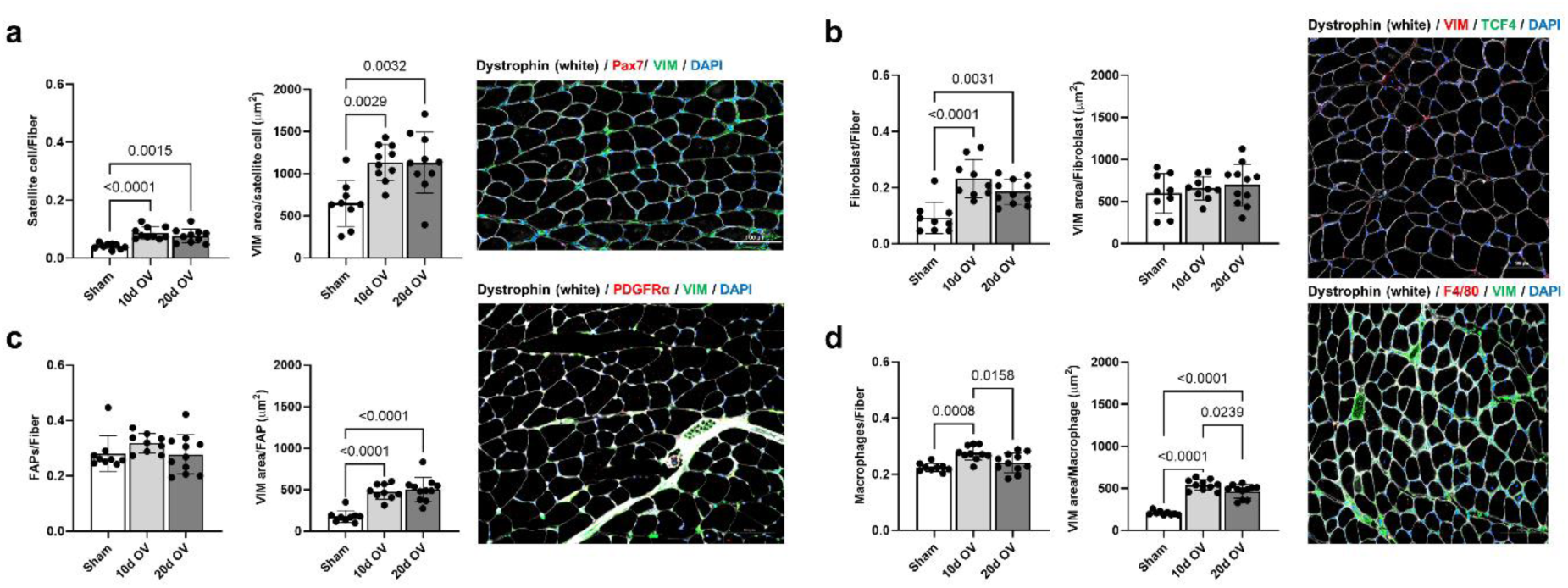
Vimentin expression with stromal cell expansion following mechanical overload. All bar graph data are presented as mean and standard deviation values with 8-11 mice per condition. a) satellite cell per fiber and VIM area per satellite cell following 10- and 20-days of MOV. b) fibroblast number per fiber and VIM area per fibroblast following 10- and 20-days of MOV. c) fibo-adipogenic (FAP) cell number per fiber and VIM area per FAP following 10- and 20-days of MOV. d) macrophage number per fiber and VIM area per macrophage following 10- and 20-days of MOV. Additional representative IHC images can be found in Supplemental Figure 6.

### Vimentin Expression in Response to MOV following the Depletion of Satellite Cells

Given the Tabula Muris and Figure 4 satellite cell data, we sought to explore the interaction between satellite cells and VIM expression following MOV. The Pax7-DTA mice (Pax7^CreER/+^; Rosa26^DTA/+^) without (vehicle) or with tamoxifen (TAM) was utilized to effectively deplete satellite cells prior to 10- and 20-days of MOV and Sham; note, general muscle characteristics are presented in Supplemental Figure 3. Satellite cells were significantly depleted in TAM-treated mice at all time points (p < 0.001; Figure 5c). VIM expression based on percent area occupied by VIM (Figure 5a) and the percent area of VIM per fiber (Figure 5b) revealed a significant treatment (TAM vs. vehicle), time, and interaction effects (all p < 0.001). Further analysis showed that VIM (% area and %area per fiber) was again responsive to 10- and 20-day MOV in vehicle-treated mice (Figure 5a/b). However, this response was significantly attenuated in TAM mice that lacked satellite cells.

**Figure 5.**
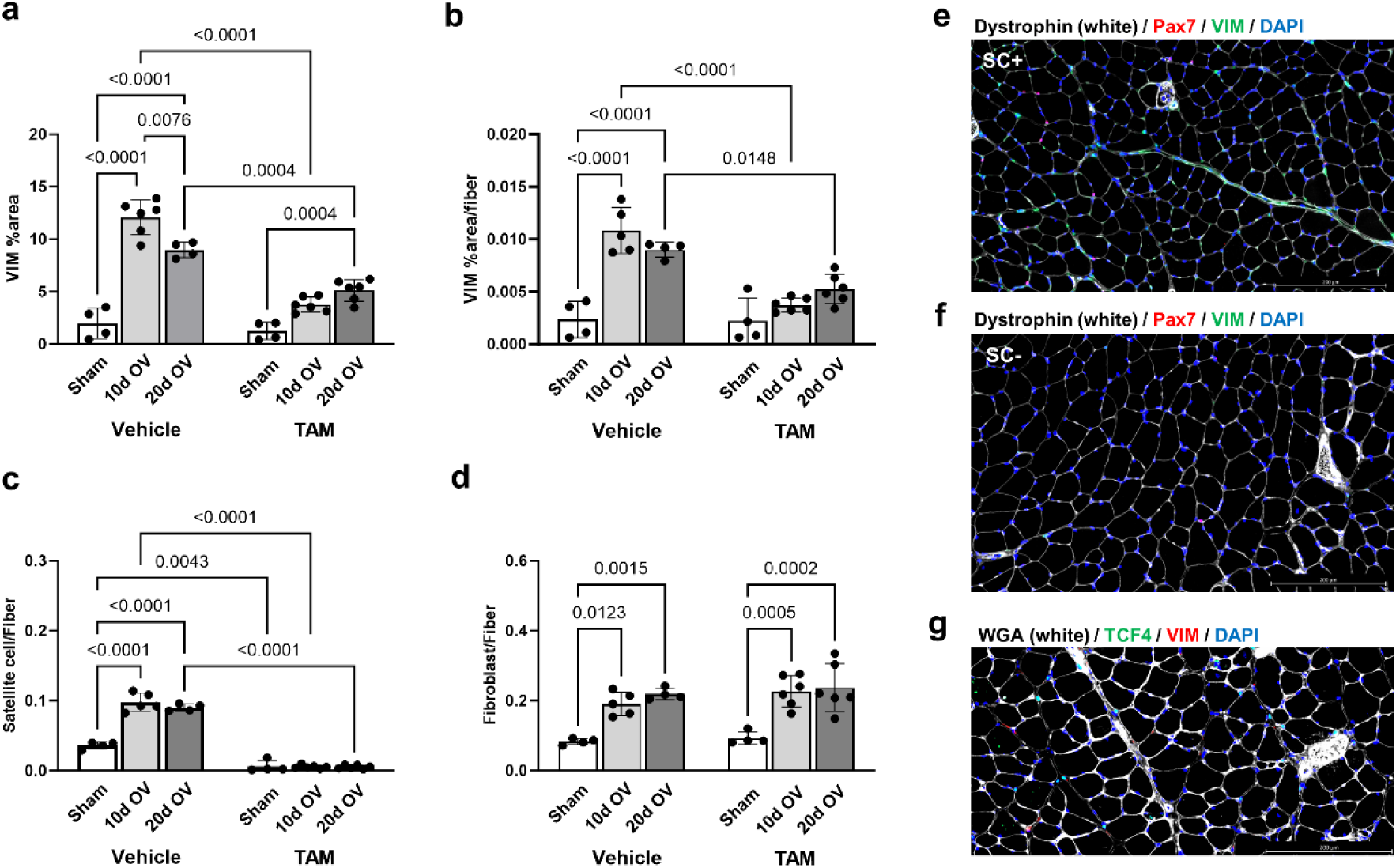
Vimentin expression in satellite cell depleted muscle following mechanical overload. All bar graph data are presented as mean and standard deviation values with 4-6 mice per condition. a) Percent area of plantaris cross-section occupied by VIM following 10- and 20- days of MOV. b) Percent area of plantaris cross-section occupied by VIM (normalized to fiber number) following 10- and 20-days of MOV. c) Satellite cell number per fiber following 10- and 20-days of MOV. d) Fibroblast number per fiber following 10- and 20-days of MOV. e) Representative image of a satellite cell intact (SC+) mouse with VIM expression. f) Representative image of a satellite cell depleted (SC-) mouse with VIM expression. g) Representative image of fibroblast staining. General model characteristics including body mass, plantaris mass, plantaris fCSA and fiber number, and myonuclei/fiber can be found in Supplemental Figure 3. Additional representative IHC images can be found in Supplemental Figure 6.

Given the modest (but significant) elevation in VIM expression following 20-days MOV in TAM-treated mice, we next measured fibroblast numbers to determine whether an increase was a potential cause for increased VIM expression. Notably, the fibroblast number responses with MOV were similar in vehicle- and TAM-treated mice (Figure 5d). Hence, we interpret these data to suggest that higher satellite cell (but not fibroblast) counts with MOV are partially responsible for the MOV-induced increases in VIM expression.

To further elucidate as to whether satellite cells are involved in VIM expression, a series of cell culture experiments were performed. In one experiment, C_2_C_12_ myotubes were treated with (AraC+) to reduce residual myoblasts. AraC+ and AraC- cells were then treated with incremental doses of rIGF-1 (as described in the Methods section). Although AraC+ and AraC- myotubes exhibited dose-dependent increases in cell diameters (Supplemental Figure 4a), VIM protein expression only increased in myotubes containing residual myoblasts (Supplemental Figure 4b). Moreover, in a separate experiment examining VIM expression during the time course of differentiation, a steady and consistent loss of VIM was observed across the 0-7-day post-differentiation timepoints (Supplemental Figure 4c). Hence, these experiments provide added confidence that satellite cells may be source of VIM. Furthermore, these *in vitro* data support that VIM expression is responsive to an anabolic stimulus.

### AAV VIM shRNA Against Vimentin Does Not Affect Measures of Gross Muscle Hypertrophy but May Lead to Delayed Long-Term Adaptations

To determine whether VIM is needed for rapid skeletal muscle hypertrophy, we used an AAV9 VIM shRNA (versus scrambled) construct to knock down VIM expression prior to 10- and 20-days of MOV. Immunohistochemistry of GFP indicated that ∼50% of the tissue in both injection groups were infected with AAV9 (Supplemental Figure 5). Plantaris masses normalized to body weights revealed a significant effect of time (p = 0.001), while no effect of treatment or interaction was detected (p = 0.887 and p = 0.334, respectively; Figure 6a). Similar outcomes were also observed for fiber number (Figure 6b), whole-muscle cross-sectional area (Figure 5c), and mean fiber cross-sectional area (Figure 6d) whereby each of these variables demonstrated significant time effects (p = 0.007, p = 0.013, and p = 0.018, respectively) but no treatment or interaction effects (p > 0.05 for all measures). Myonuclei per fiber revealed significant time and treatment effects (p = 0.012 and p = 0.019), but not a significant interaction (p = 0.279; Figure 6e).

**Figure 6.**
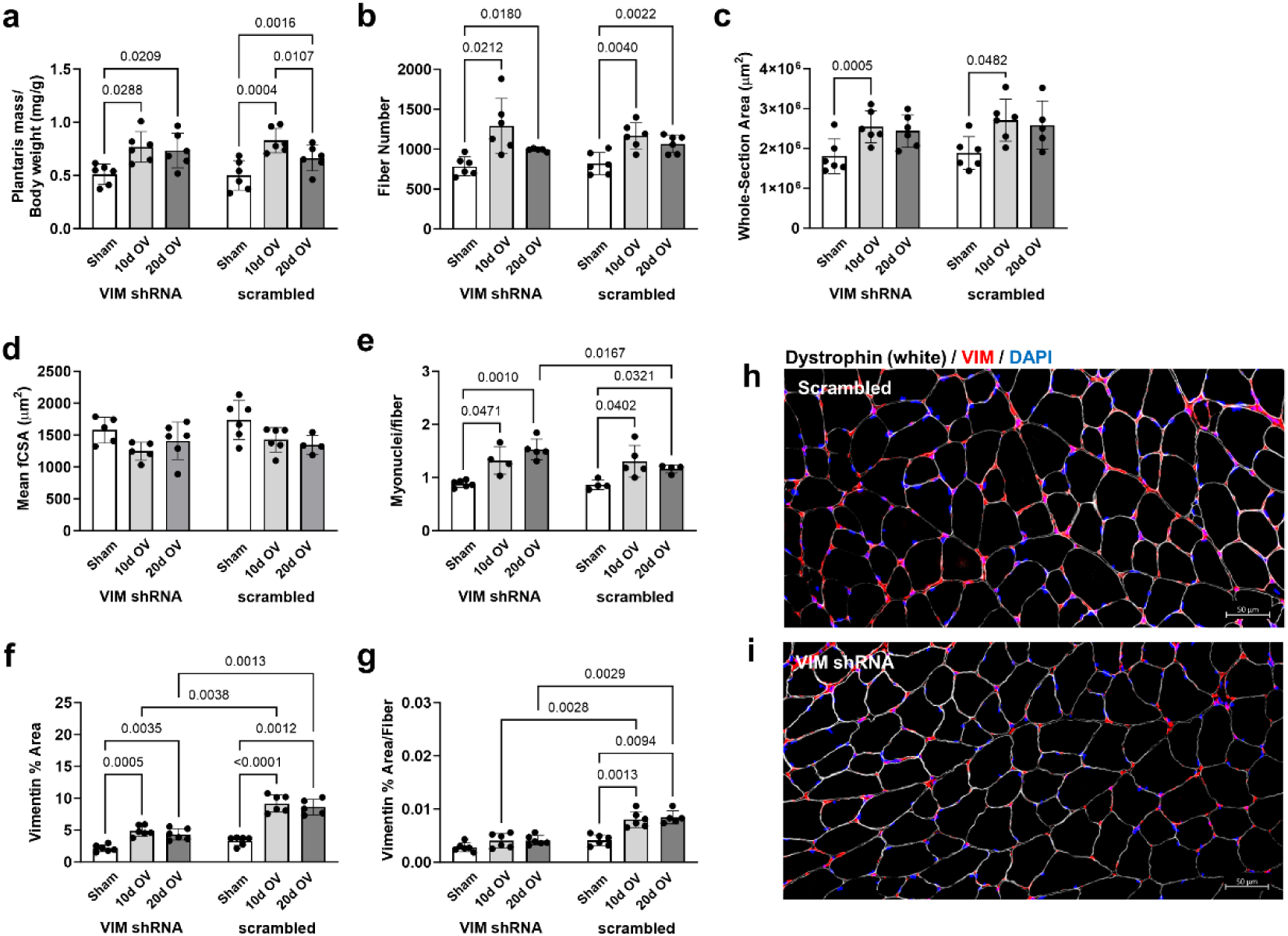
General muscle characteristics and vimentin expression in mice treated with AAV9 VIM shRNA. All bar graph data are presented as mean and standard deviation values with 4-6 mice per condition. a) Plantaris masses (normalized wet to body weight) following 10- and 20-days of MOV. b) Number of fibers present in whole cross-section following 10- and 20-days of MOV. c) Cross-sectional area of whole section following 10- and 20-days of MOV. d) mean fiber cross-sectional area following 10- and 20-days of MOV. e) Myonuclei per fiber following 10- and 20-days of MOV. f) Percent area of cross-section occupied by VIM following 10- and 20-days of MOV. g) Percent area of cross-section occupied by VIM (normalized to fiber number) following 10- and 20-days of MOV. h) Representative image for dystrophin (fCSA) and VIM abundance in scrambled mice. i) Representative image for dystrophin (fCSA) and VIM abundance in AAV-VIM shRNA mice. Additional representative IHC images can be found in Supplemental Figure 7.

VIM expression as measured by the percent of total area occupied and the percent area normalized to fiber number revealed significant time (p < 0.001 for both), treatment (p < 0.001 for both), and interaction effects (p < 0.001 and p = 0.007, respectively). Further analysis revealed that VIM expression measured as both percent area of VIM and percent area of VIM per fiber was significantly lower in the VIM shRNA mice following both 10- and 20-day MOV compared to scrambled mice (10-day p = 0.004 and p = 0.003; 20-day p = 0.001 and p = 0.003; Figure 6f/g).

While a blunting of VIM expression did not affect measures of gross skeletal muscle hypertrophy following 10- and 20-day MOV, a noted shift toward smaller fibers was evident in VIM shRNA versus scrambled mice based on fCSA histogram distributions (Figure 7a/b). In this regard, Kruskal-Wallis ANOVA with Bonferroni correction indicate that in both VIM shRNA and scrambled mice, 10- and 20-day MOV fCSA distribution was lower than Sham (p < 0.001 for all). Further, 10- and 20-day MOV VIM shRNA mice had lower fCSA distribution compared to all other groups (p < 0.001 for all), except for no difference between 20-day VIM shRNA and 20-day scrambled mice (p = 0.110). Evidence of morphologically impaired myofiber hypertrophy was also observed in VIM shRNA mice who presented more centrally located myonuclei (interaction p < 0.001; Figure 7c) and myofibers that stained positive for embryonic myosin (interaction p < 0.001; Figure 7e).

**Figure 7.**
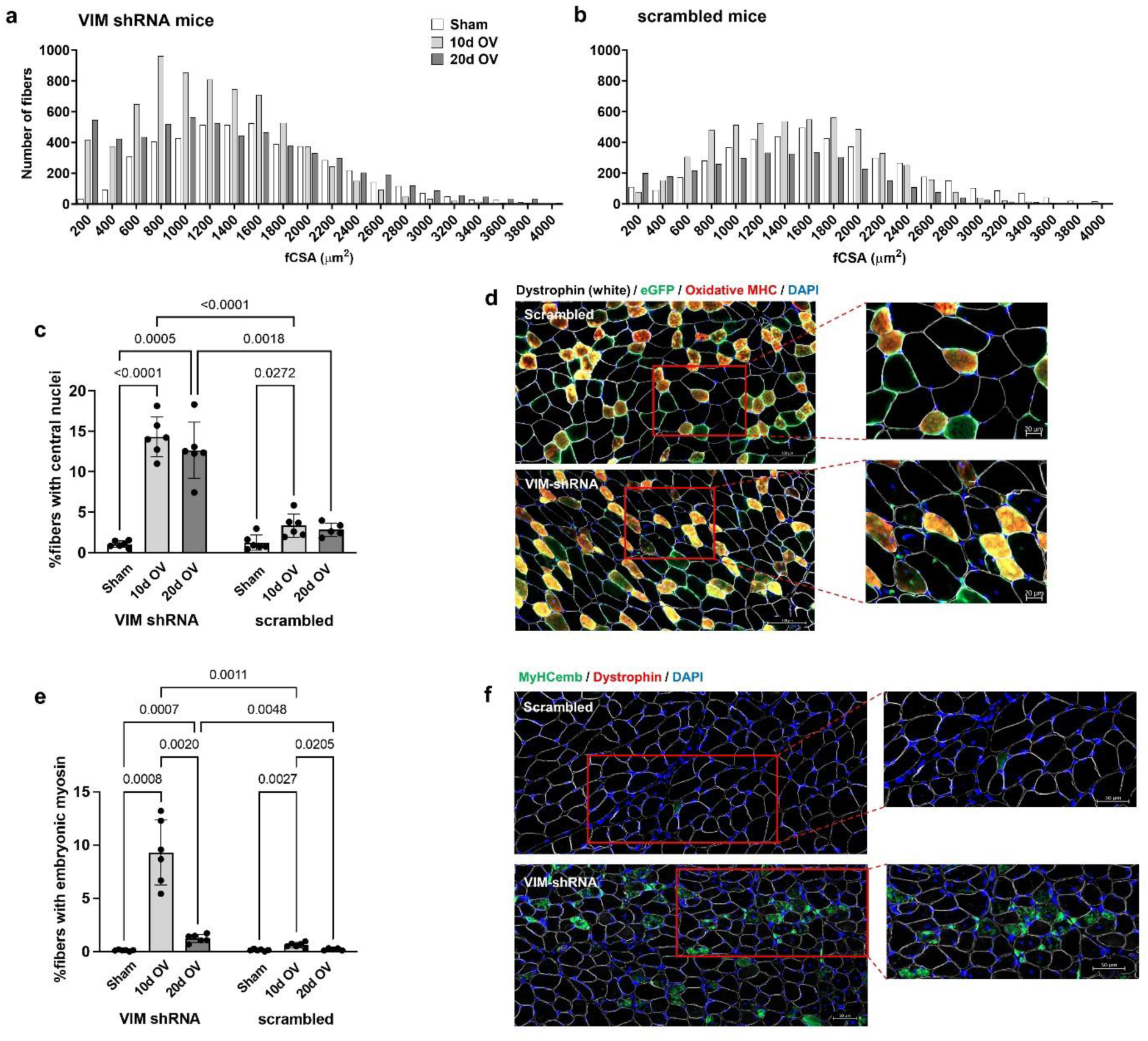
VIM shRNA mice present a shift toward smaller fibers and more centrally located nuclei. All bar graph data are presented as mean and standard deviation values with 4-6 mice per condition. a) Histogram for fiber size distribution in VIM shRNA following 10- and 20-day MOV. b) Histogram for fiber size distribution in scrambled mice following 10- and 20-day MOV. c) Percentage of fibers that display centrally located nuclei following 10- and 20-day MOV between AAV9 injection groups with representative IHC images in panel d. e) Percentage of fibers positive for embryonic myosin heavy chain (MyHC_emb_) following 10- and 20-day MOV between AAV9 injection groups with representative IHC images in panel f. Additional representative IHC images can be found in Supplemental Figure 7.

## DISCUSSION

Skeletal muscle VIM expression is responsive to mechanical loading as evidenced by increases in this protein following 10 weeks of resistance training in humans and a higher presence of this protein in the ECM following 10- and 20-days of mechanical overload in mice. Given past research linking VIM expression to satellite cells [19, 22], we used the Pax7-DTA mouse model to determine the relationship between satellite cell abundance and VIM expression. This model showed that VIM expression is lower in satellite cell-depleted mice following MOV compared to satellite cell-intact mice, and our *in vitro* data lend further potential evidence of satellite cells being a potential source of VIM. Lastly, targeted viral knockdown with an AAV9 VIM shRNA resulted in a blunting of VIM expression following 10- and 20-days of MOV in mice compared to scrambled mice. This blunting of VIM expression did not affect gross measures of skeletal muscle growth (i.e., plantaris weight, fiber number, mean fCSA). However, a notable increase in the number of smaller fibers was evident in VIM shRNA mice and this was accompanied by a marked increase in the number of MyHC_emb_-positive fibers with centrally located nuclei following 10- and 20-days of MOV. A summary of these findings is presented in Figure 8.

**Figure 8.**
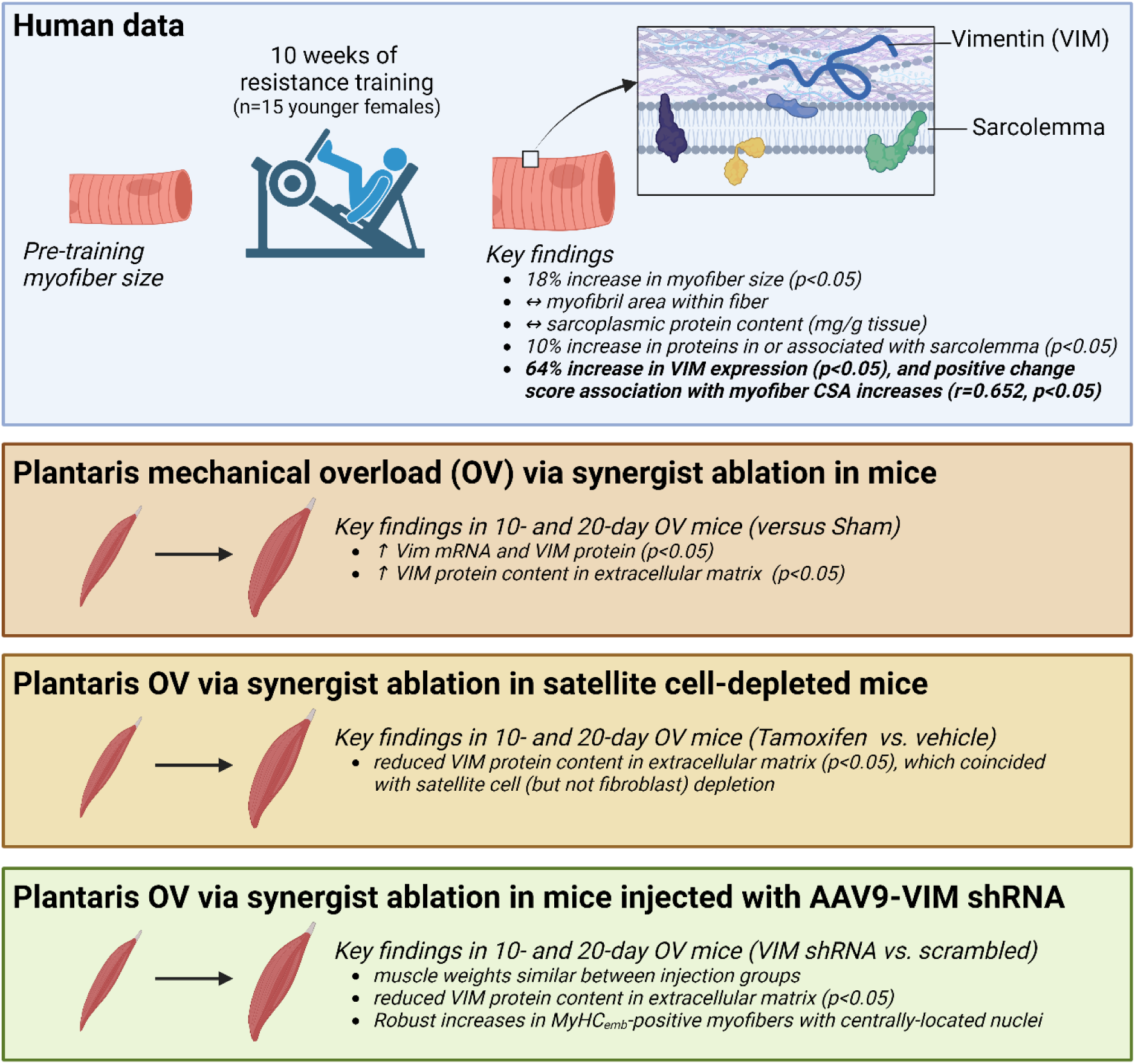
Summary of findings. Figure illustrates summary of findings in humans and mice. Schematic created using a BioRender site license subscription.

As no other study has sought to directly examine how VIM is related to skeletal muscle hypertrophy, our discussion is somewhat limited to hypotheses based on our observations. However, there are data consistent with some of our findings. Using a time course of MOV in mice, Chaillou et al. 2013 [7] reported that *Vim* mRNA expression is elevated in skeletal muscle and further links VIM as a potential mechanosensitive gene in skeletal muscle upregulated in response to synergist ablation. Moreover, the knockdown or knockout of VIM expression has been linked to decreases in cell size and protein synthesis pathways [25, 37–39], albeit these studies have examined other cell types and our fCSA and whole section CSA data are conceptually discordant with these reports. Perhaps what is consistent with our findings, however, are select studies indicating that VIM is involved with regeneration in several tissues including skeletal muscle [23, 40]. In this regard, VIM expression during regeneration in skeletal muscle has been shown to be elevated 7-days post injury, and others have reported that VIM expression is linked to key aspects of satellite cell involvement in the regeneration process [19, 22, 41]. Hence, these prior reports along with our observations of VIM knockdown promoting a regenerative phenotype following MOV warrant continued research in this area of muscle biology.

Aside from the current observations, several outstanding issues remain. First, while we posit that VIM is a mechanosensitive target, our *in vitro* data indicate that this target is also responsive to IGF-1. Indeed, both stimuli elicit similar cell signaling events (e.g., mammalian target of rapamycin complex-1 activation), and there are distinct differences between growth factor signaling and mechanotransduction in skeletal muscle [1]. Also notable are studies that have mapped the VIM promoter and suggested that various transcription factors driven by pro-inflammatory ligands (e.g., heterodimers binding to AP-1 sites, NF-κB) drive gene expression [42, 43]. Hence, investigations parsing out the upstream signals that upregulate VIM gene/protein expression in skeletal muscle during rapid growth are needed.

### Limitations

This study has various limitations. First, the bulk of our investigation hinged on rodent findings, and this was due to human tissue limitations. Hence, future time course resistance training investigations should aim to implement IHC to replicate our rodent findings. The C2C12 in vitro experiments, while insightful, do not represent satellite cell dynamics given that they are an immortalized cell line. Another limitation is that the AAV9 experiments utilized unilateral synergist ablation and the growth response compared to the dual leg synergist ablation model utilized for the other mouse experiments in this study resulted in differential hypertrophy responses (see fCSA profiles of Figure 3e versus 7b). We also did not examine whether shRNA-mediated VIM knockdown impaired muscle function and this requires further inquiry. Finally, while the shRNA experiments performed herein provide valuable preliminary evidence in this area of muscle biology, elegant genetic mouse models knocking out VIM in a cell-specific fashion (e.g., satellite cell VIM-KO mice) could continue to confirm or refute our hypotheses put forth herein.

### Conclusions

Using an integrative approach involving cell culture, rodent, and human experiments, we posit that VIM is a mechanosensitive gene in skeletal muscle that is predominantly localized to the ECM. Moreover, the disruption of VIM expression during periods of mechanical overload leads to a dysregulated regenerative response that is suggestive of dysfunctional skeletal muscle hypertrophy.

## AUTHOR CONTRIBUTIONS

This study fulfilled the dissertation requirements for J.S.G., and J.S.G., C.B.M., and M.D.R. conceived the idea for this study. C.S.F., A.D.F and A.N.K. were committee members who provided continued support throughout the project. C.A.L., I.V., A.D.F., J.J.M., and M.D.R. provided significant resources to complete various project aims. J.M.M. and M.M. performed multiple experiments throughout the study, which warranted co-authorship. J.S.G. and M.D.R. primarily drafted the manuscript, and all co-authors provided feedback as well as intellectual contributions. All authors have read and approved the final version of this manuscript and agree to be accountable for all aspects of the work in ensuring that questions related to the accuracy or integrity of any part of the work are appropriately investigated and resolved. All persons designated as authors qualify for authorship.

## ACKNOWLEDGMENTS

Reagent costs (e.g., reagents, antibodies, and mice) were purchased through discretionary laboratory funds of M.D.R., I.V., and C.A.L. C.A.L. was also supported by the São Paulo Research Foundation (#2020/13613-4) and the National Council for Scientific and Technological Development (#311387/2021-7).

## DATA AVAILABILITY STATEMENT

The data that support the findings of this study are available from the corresponding author upon reasonable request.

