## Supplemental figures for "Resistance Exercise and Mechanical Overload Upregulate Vimentin for Skeletal Muscle Remodeling"

**Supplemental Figure 1.** General muscle characteristics from human participants following 10-weeks of resistance training.

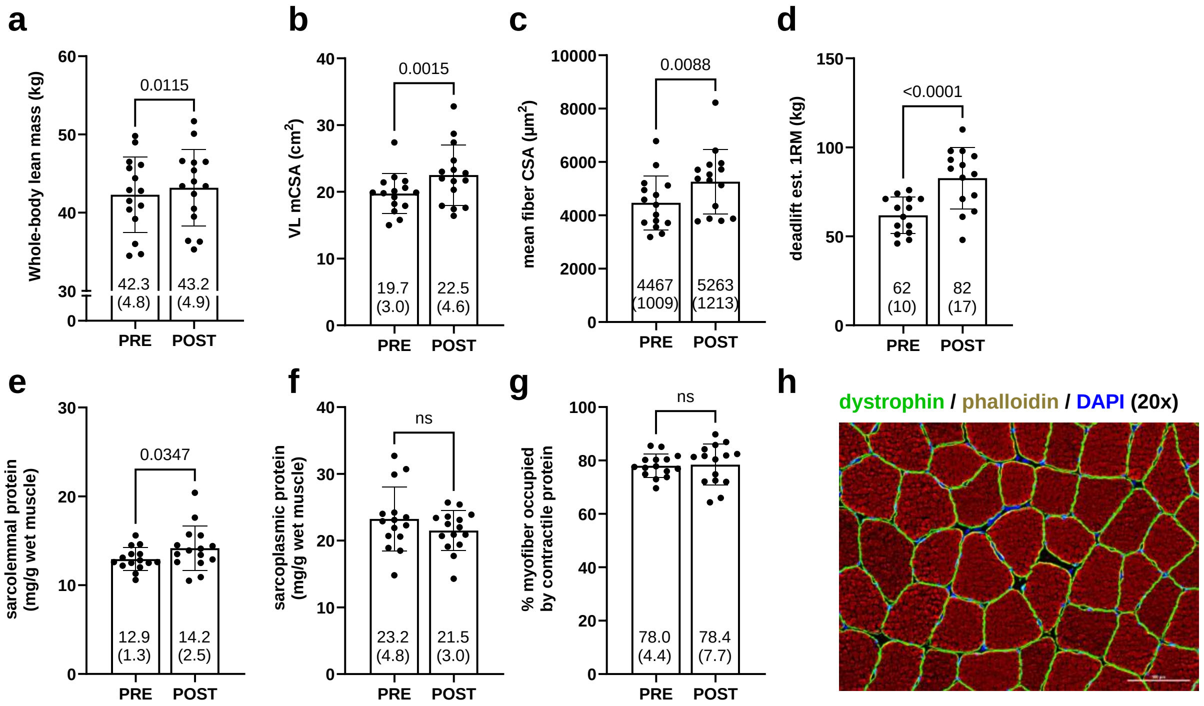


Legend: a) whole body lean mass measured via DEXA. b) vastus lateralis muscle cross-sectional area measured via panoramic ultrasound. c) mean fiber cross-sectional area measured via IHC. d) deadlift 1RM estimated from 3RM maximum testing. e) sarcolemmal protein concentrations. f) sarcoplasmic protein concentrations. g) percent area of myofibers occupied by myofibrils measured via phalloidin staining and IHC. h) Representative image of phalloidin staining used to measure myofibril content. Additional note: data contain 15 participants for all outcomes.

**Supplemental Figure 2.** General characteristics following 10- and 20-days of plantaris overload in wild type mice (accompanies Fig. 3 data).

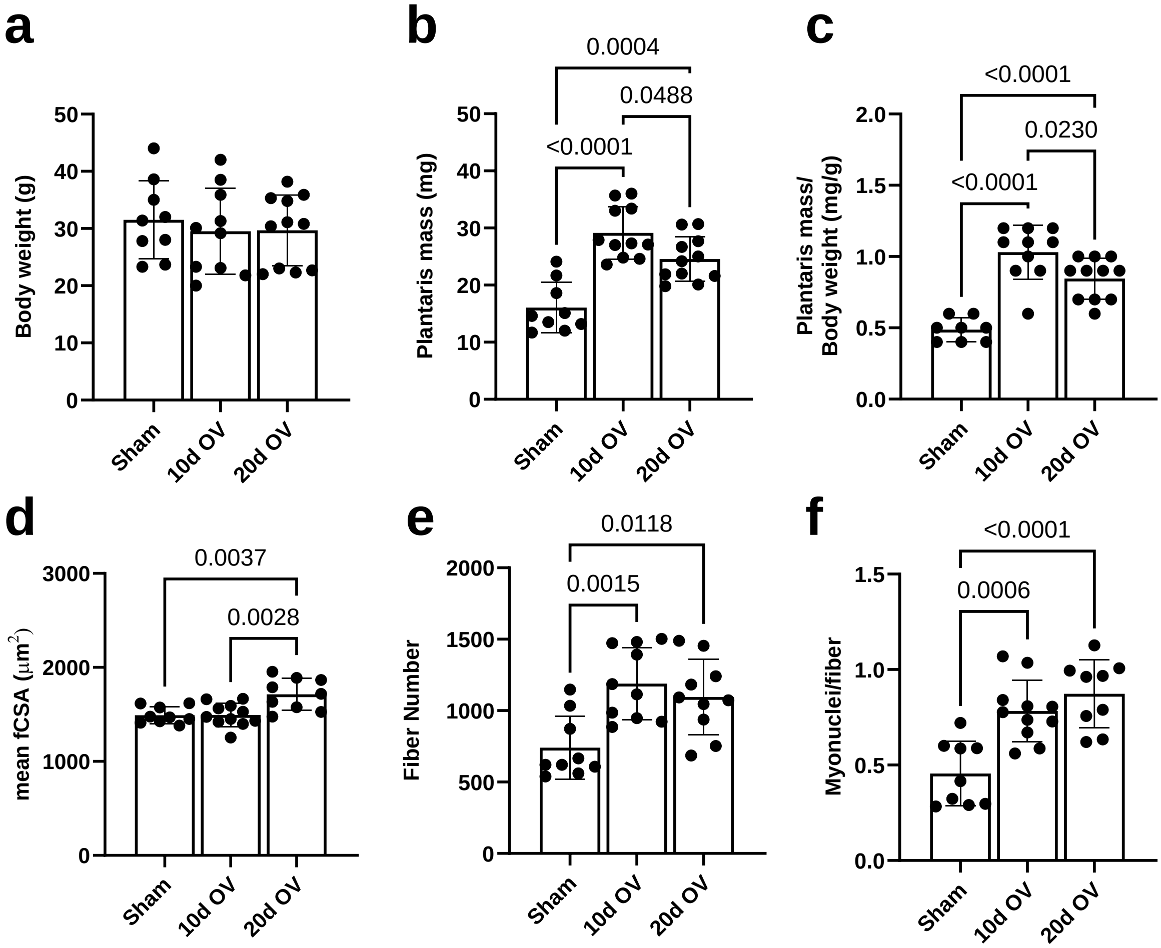


Legend: a-c) Body weight, plantaris mass, and plantaris weight normalized to body weight. d) mean fiber cross-sectional area measured via IHC. e) total number of fiber counted from whole cross-section. f) Myonuclear number per fiber measured from cross-section. Additional note: bar graphs are presented as mean and standard deviation values and data contain 8-11 mice per condition.

**Supplemental Figure 3**. General Muscle Characteristics of Pax7-DTA Mice (accompanies Fig. 5 data).

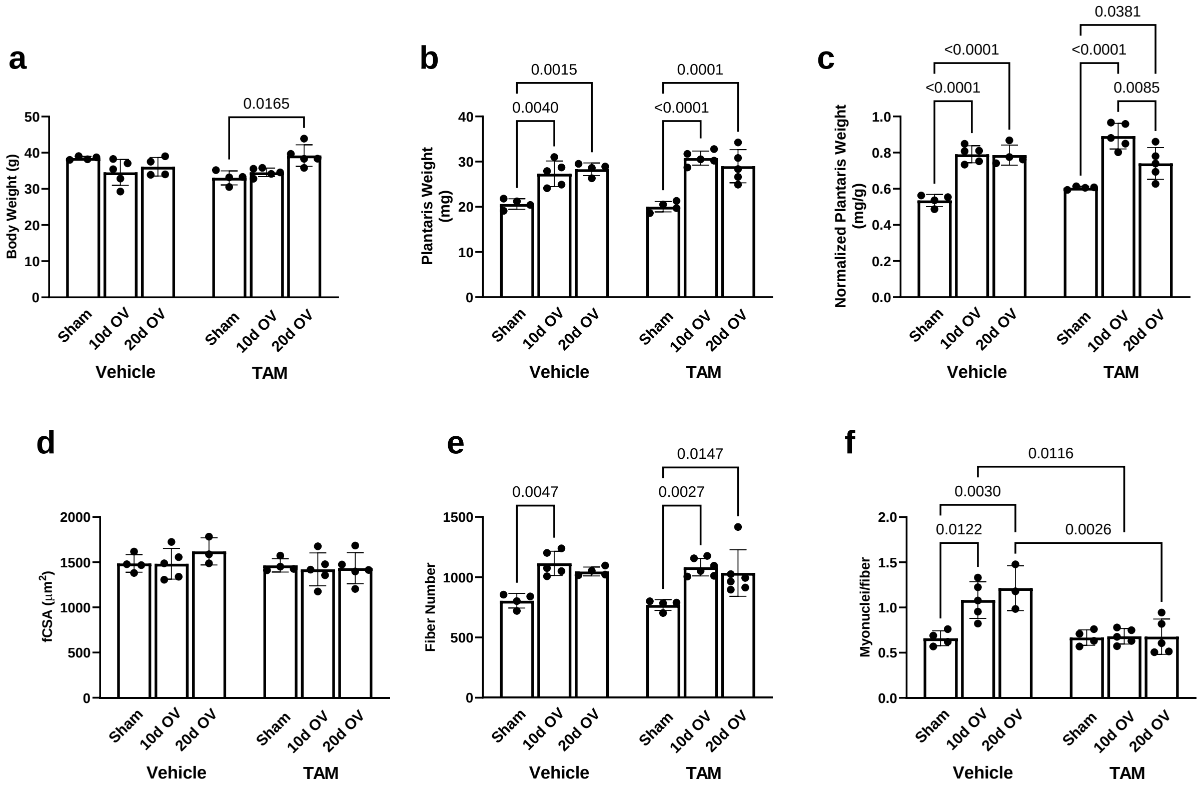


Legend: a-c) Body weight, plantaris mass, and plantaris weight normalized to body weight. d) mean fiber cross-sectional area measured via IHC. e) total number of fibers counted from whole cross-section. f) Myonuclear number per fiber measured from cross-section. Additional note: bar graphs are presented as mean and standard deviation values, data contain 4-5 mice per condition.

**Supplemental Figure 4.** Vimentin expression in $C_2C_{12}$ myotubes during differentiation and with an anabolic stimulus.

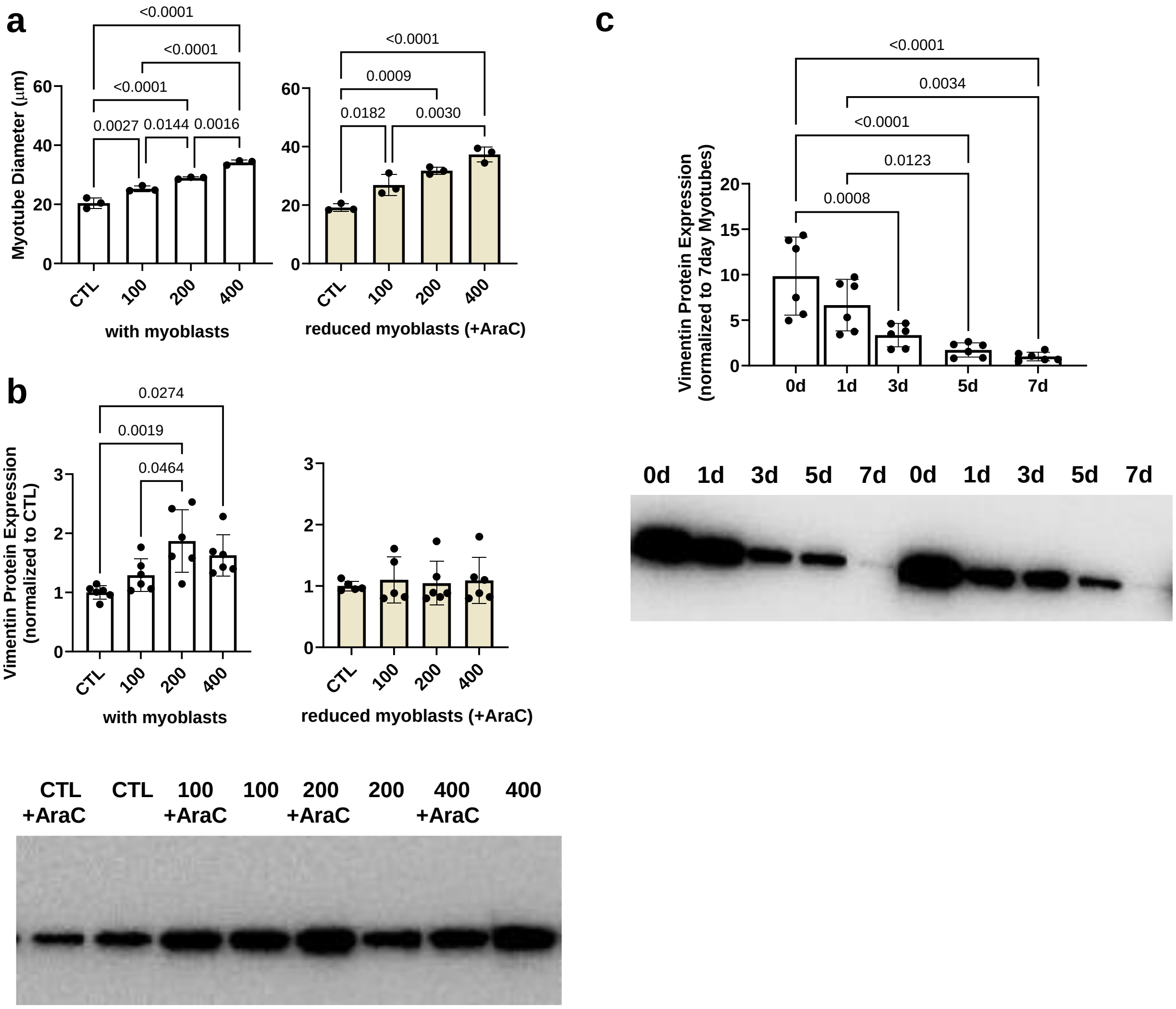


Legend: a) Myotube diameters of $C_2C_{12}$ following incremental 24-hour dosages of mouse rIGF-1 (ng/mL) with residual myoblasts left intact or reduced in number via 24-hour pre-treatments of 10 µM arabinosylcytosine (AraC). b) VIM protein expression these same experiments with representative western blot images below. c) VIM protein expression during a time course of various stages of differentiation in $C_2C_{12}$ myoblasts with representative western blot images below.

**Supplemental Figure 5.** GFP-positive myofibers in AAV9-injected mouse plantaris muscles (accompanies Fig. 6/7 data).

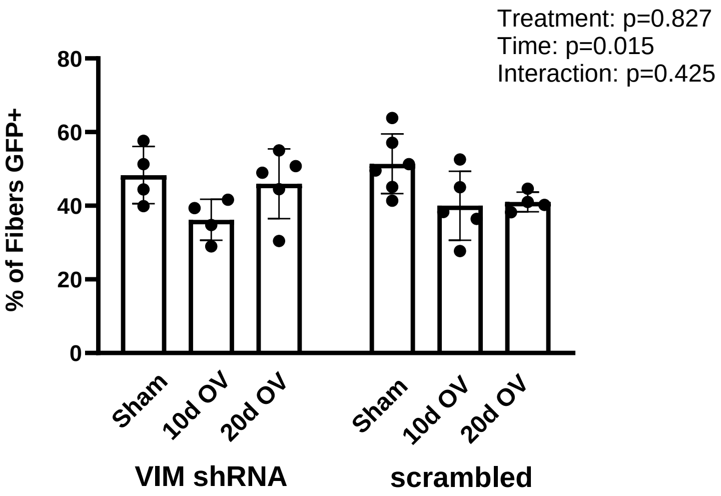


Legend: Data show ~50% successful viral transduction (grand mean) in both injection groups for the experiment.

**Supplemental Figure 6.** Additional representative images (accompanies Fig. 3/4/5 data).

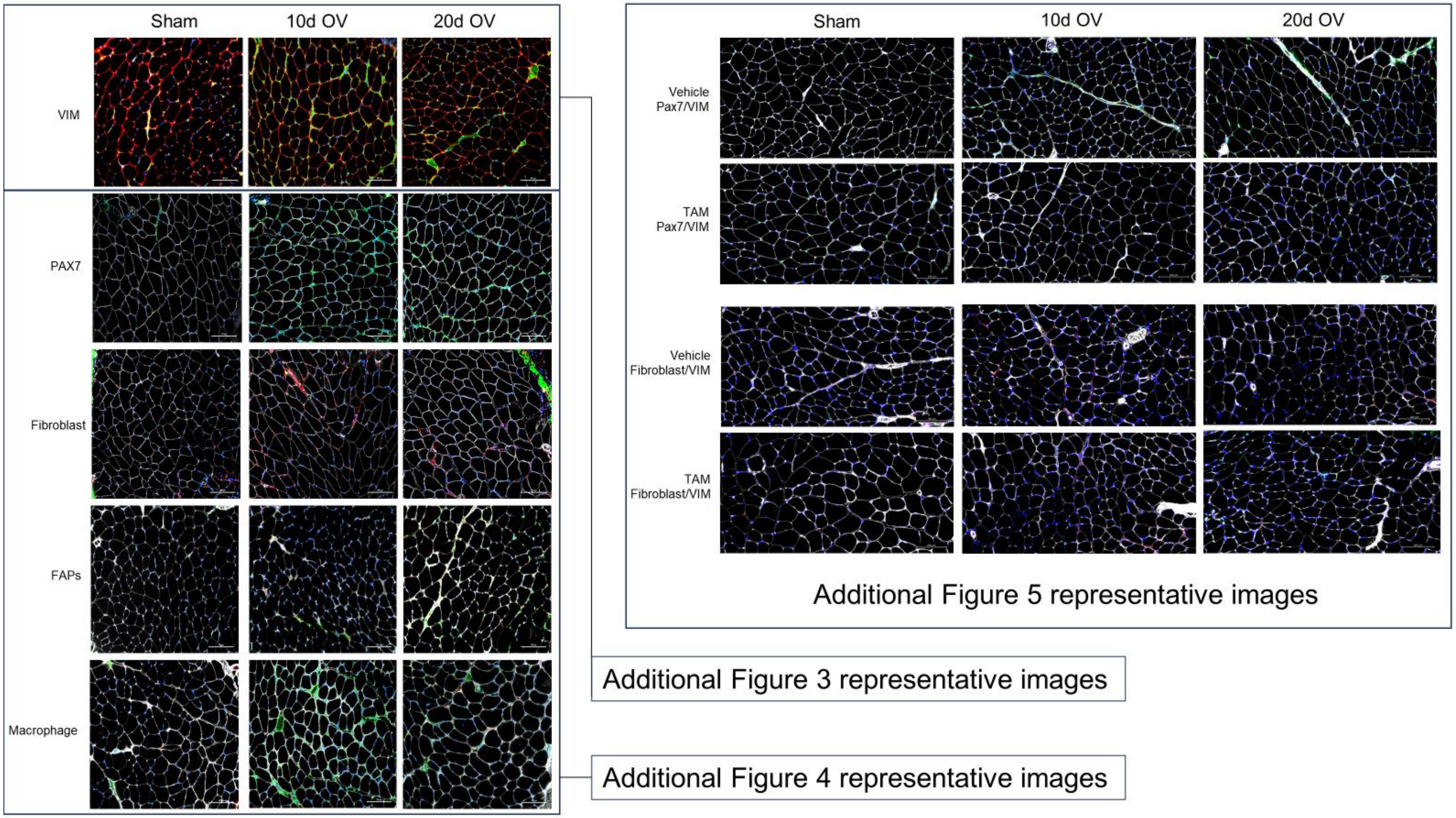

**Supplemental Figure 7.** Additional representative images (accompanies Fig. 6/7 data).

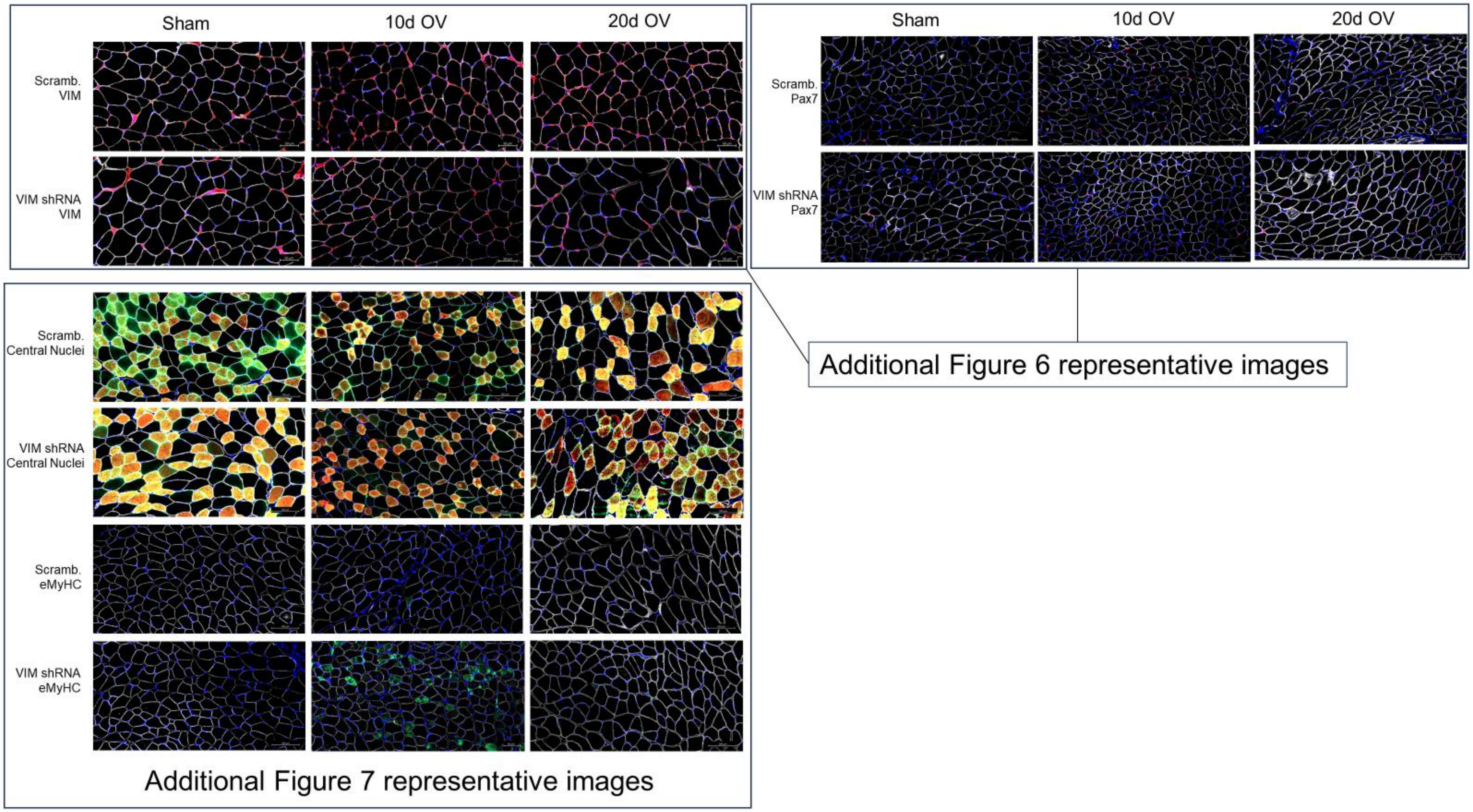
